# Manganese Accumulation for Genetically Induced Contrast (MAGIC) MRI in the brain across species

**DOI:** 10.64898/2026.04.30.722028

**Authors:** Harikrishna Rallapalli, Dwina Hadj-Mabrouk, Tyler Lee, Wenliang Wang, Walter Lerchner, Keith MacRenaris, Tom O’Halloran, Barry Richmond, Alan Koretsky

## Abstract

Mapping the mesoscale architecture of neural circuits is essential for understanding brain function, yet high-resolution anatomical tracing remains largely dependent on fluorescent reporters that require terminal histology. Here, we present Manganese Accumulation for Genetically Inducible Contrast (MAGIC) MRI, a gene expression reporter system based on the metal ion transporter Zip14 (Slc39a14) which enables noninvasive, in vivo neural tracing. In rodents, viral delivery of Zip14 enables both anterograde and retrograde tracing of cortico-thalamic and basal ganglia circuits. Mechanistic validation via laser ablation-inductively coupled plasma-time-of-flight-mass spectrometry (LA-ICP-TOF-MS) confirmed that MRI contrast changes are driven by specific Mn^2+^ accumulation. This permitted high-resolution visualization of neural populations and projections using clinical standard MRI sequences without supplementary contrast agents. However, addition of systemic Mn^2+^ further increased the signal by a factor of 2-5 fold. To facilitate objective, high-throughput analysis, we developed a fully automated pipeline for voxel-wise anomaly detection that accurately identifies and quantifies MAGIC enhanced regions in individual subjects. Finally, it is demonstrated that MAGIC is translatable to the large mammalian brain, providing the first functional demonstration of an MRI-visible reporter in the rhesus macaque. By enabling the non-invasive monitoring of neural connectivity across species, MAGIC provides a versatile, longitudinal tool for studying structural plasticity and circuit organization in the living brain.

## Main

Genetically encoded reporters visible using MRI (GEMRI) promise a high-resolution, noninvasive readout that maps gene expression to mesoscopic anatomy without optics, implants, or ionizing radiation^1^. Yet despite three decades of innovation, most GEMRI strategies remain at the proof-of-principle stage. Key barriers include insufficient sensitivity at clinical field strengths, reliance on bespoke pulse sequences or specialized chemistries, and reliance on tissue-targeted delivery of exogenous agents. Furthermore, none have yet demonstrated broad utilization or efficacy in species other than rodents. A reporter that uses standard acquisition protocols, leverages endogenous metabolism, and proves effective in different species, especially the old-world primate brain, would address these challenges and open a practical route toward clinical application of GEMRI.

Here we extend development of such a platform using the metal transporter Zip14 (Slc39a14) as a GEMRI reporter we refer to as Manganese Accumulation for Genetically Inducible Contrast (MAGIC) MRI. Zip14 expression can concentrate endogenous metals, predominantly Mn^2+^, within cells to produce T_1_ shortening detectable with off-the-shelf MRI pulse sequences. We have demonstrated that expressing Zip14 using well-characterized adeno-associated and lentiviral capsids can enable MRI to read out anterograde and retrograde projections in the rodent brain, yielding layer-resolved cortical-thalamic and striatal-pallidal-nigral maps in vivo^2^. We confirm that the mechanism of the enhancement is specifically due to increased Mn^2+^ in areas expressing Zip14. MAGIC MRI can operate without exogenous agents. However, MAGIC MRI can be amplified upwards of two to five-fold using well-established Mn^2+^ supplementation procedures (MAGIC+), enabling automated detection of Zip14 related enhancement over the whole mouse brain and increased sensitivity to neural connectivity. Critically, the successful application of MAGIC in the rhesus macaque brain is demonstrated using a standard clinical 3 T MRI. Zip14 is the first GEMRI agent to cross the primate translational barrier, establishing a powerful and clinically relevant platform for studying large-scale neural circuits in higher-order species.

## Results

### MAGIC enables in vivo mesoscale neural tracing

To prepare for mechanistic analysis of MAGIC and to determine the benefits of MAGIC+, the metal ion transporter Zip14 was expressed in the mouse brain to quantify the signal changes obtained (Fig 1 a). Plasmid constructs consisting of the neuronal-specific Synapsin promoter (Syn), Slc39a14 gene (coding for Zip14), and a histology-visible tag (EGFP or FLAG) were packaged into AAV9 or RetroAAV9 as previously described^2^. These viruses were stereotactically injected into the mouse brain at coordinates targeting the primary somatosensory barrel cortex (S1BC) or the caudate putamen (CP) in the right hemisphere. These mice were imaged using an 11.7 T small animal MRI system one week before (Pre) and two weeks after AAV delivery using the vendor-available Magnetization Preparation Rapid Gradient Echo (MPRAGE) T_1_ weighted imaging protocol.

**Fig 1.**
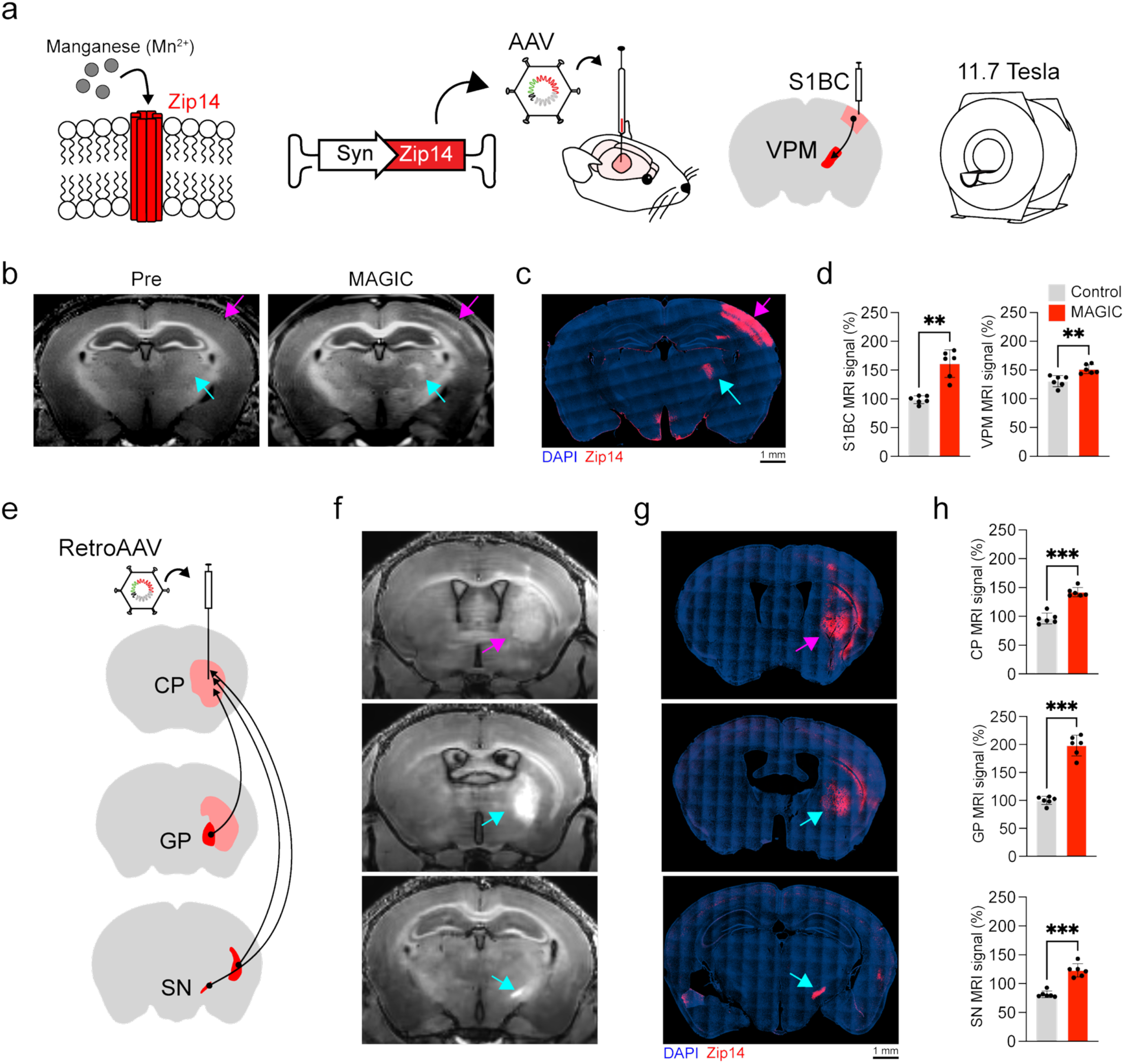
MAGIC enables in vivo mesoscale neural tracing. (a) The manganese transporter Zip14 was expressed in the mouse brain using transcranially delivered AAVs under the control of the neuronal specific Synapsin (Syn) promoter. Imaging was performed before and at several timepoints after injection using an 11.7 T small animal MRI system. (b) In vivo MAGIC in the injected S1BC (magenta arrow) and primary anterograde projection site, the ipsilateral VPM (cyan arrow), was detected that was not observed on the Pre scan (n=6). (c) This enhancement closely correlates with ex vivo immunohistology for Zip14. (d) These effects were reproducible across animals and statistically significant. (e,f) In vivo retrograde tracing of basal ganglia circuits was potentiated by RetroAAV-Zip14 injection into the caudate putamen (CP) which receives projections from neurons housed in the globus pallidus (GP) and substantia nigra (SN) (n=6). (g) Bright MAGIC was apparent in all three areas and well-correlated with Zip14 immunohistology. (h) These effects were reproducible across animals and statistically significant. P-value thresholds: ^*^p<0.05 ^* *^p<0.01 ^***^ p<0.001.

As expected, Pre scan images provided contrast between the cortical gray matter and relatively hyperintense white matter and hippocampus (Fig 1 b). AAV-Zip14 injections into the S1BC produced focal hyperintensity on MAGIC images at the injection site and projecting areas (Fig 1 b). Specifically enhanced cortical layers included I, II/III, and V and was consistent with our previous application of MAGIC in the rodent S1BC^2^. Bright contrast was also focally apparent in the VPM ipsilateral to the injected S1BC, in addition to more widespread, moderate hyperintensity in the other areas known to contain projecting axons from the neurons in the S1BC to the VPM. Matched immunofluorescence histology for Zip14 confirmed that the observed MAGIC enhancement was colocalized with increased expression of Zip14 (Fig 1 c). This demonstrated that MAGIC MRI enables anterograde neural tracing. Staining for the neuronal marker NeuN confirmed that increased Zip14 expression was produced by AAV-Zip14 transduced neurons (Supplementary Figure 1). MAGIC MRI in the injected S1BC and VPM was robust and reproducible (Supplementary Table 1, Fig 1 d).

To determine which brain areas were best for quantitative metal concentration analyses, RetroAAV-Zip14 was injected into the CP of the mouse brain (Fig 1 e). On MAGIC images, bright contrast enhancement was apparent in the injected CP, and in basal ganglia areas containing neurons which project to the CP. These include the globus pallidus (GP) and substantia nigra (SN). Both the GP and the SN were focally hyperintense compared to the same areas in the contralateral hemisphere indicating that RetroAAV enabled MAGIC MRI to trace retrograde connections (Fig 1 f). Matched histology confirmed that the MAGIC hyperintensity was produced by Zip14 expression in the CP, GP, and SN (Fig 1 g). Quantification of the MAGIC changes in all areas across subjects showed robust and reproducible effects (Supplementary Table 1, Fig 1 h).

### MAGIC is produced by accumulation of endogenous Mn^2+^

Zip14 is an essential metal ion transporter which is known to transport Mn^2+^, Fe^2+^, Cu^2+^, and Zn^2+^. Mn^2+^, Fe^2+^, and Cu^2+^ are paramagnetic and thus can act as MRI contrast agents with differing magnetization relaxation properties. Quantification of Zip14 metal transport has not been performed in vivo, but mutations in Zip14 in humans and mice lead to alterations in Mn^2+^ homeostasis with much smaller effects on Fe^2+^ and Zn^2+ 3,4^. To determine the fate of metal ion levels in Zip14 expressing areas, laser ablation inductively coupled plasma time-of-flight mass spectrometry (LA-ICP-TOF-MS) was used which provides spatially resolved full mass spectrum data on metals at resolutions higher than, but comparable to, MRI. Mice were injected with RetroAAV-Zip14 into the CP due to the large MAGIC MRI enhanced area in multiple brain areas. MRI was performed before and at least two weeks after RetroAAV-Zip14 injection using the standard MPRAGE protocol. Quantitative R_1_ (80 µm in plane) and R_2_* MRI (80 µm isotropic) maps were made. After the final imaging session, 20 µm thick snap frozen brain tissue sections at the level of the CP, GP, and SN were analyzed using LA-ICP-TOF-MS at 20 µm spot size to measure in situ metal concentrations (Supplementary Figure 2, Fig 2 a).

**Fig 2.**
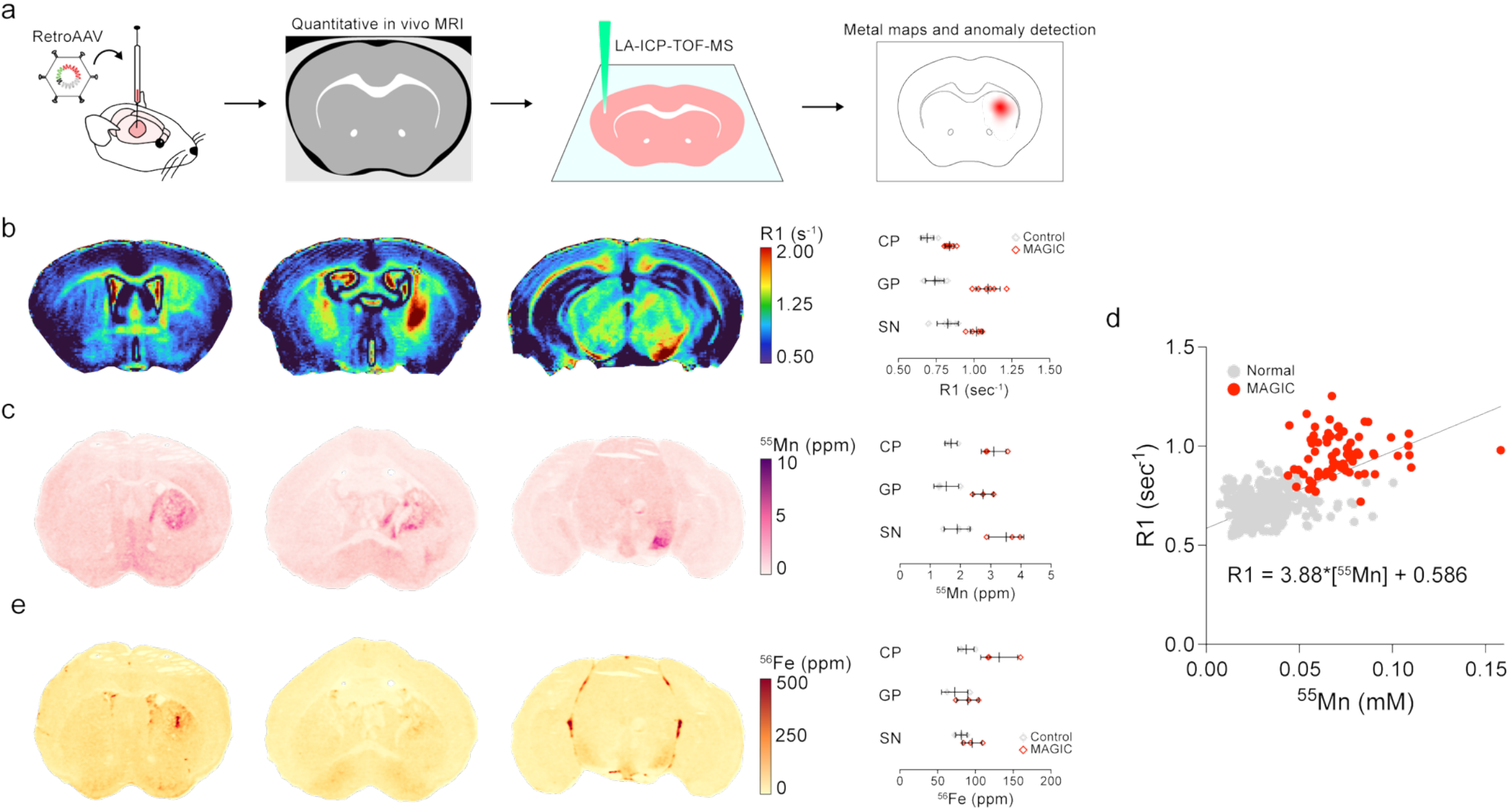
MAGIC is produced by accumulation of endogenous Mn^2+^. (a) RetroAAV-Zip14 was injected into the CP for in vivo retrograde tracing using MAGIC and quantitative R_1_ and R_2_* mapping was performed. After the last imaging timepoint, animals were sacrificed, brains were extracted, cryosectioned, and LA-ICP-TOF-MS was performed to obtain in situ metal concentration measurements. Metal maps and quantitative MRI maps were correlated. (b) Representative R_1_ mapping results reveal observable R_1_ increases in the CP, GP, and SN. ROI-based quantification of this effect confirmed that the R_1_ increases were statistically significant and reproducible across animals (n=6). (c) ^55^Mn concentration maps demonstrated Zip14 expressing areas in the CP, GP, and SN had markedly increased levels compared to endogenous levels ^55^Mn. Automated ROI-based quantification of ^55^Mn in these areas confirmed Zip14-expressing areas had significantly increased ^55^Mn, up to double the endogenous level on average (n=3). (d) Matched ROI measurements (n=450, 320 µm diameter circles) made on R_1_ maps and ^55^Mn maps enabled measurement of in vivo longitudinal relaxivity for Mn^2+^. (e) ^56^Fe concentration maps revealed increased levels in the CP, but no other brain area. Automated ROI-based quantification of ^56^Fe in these areas revealed significant increases in the CP and SN, but not the GP (n=3). P-value thresholds: ^*^p<0.05 ^* *^p<0.01 ^* * *^p<0.001

R_1_ mapping of RetroAAV-Zip14 injected mice revealed robust and reproducible increases in R_1_ in the injected CP and ipsilateral GP and SN compared to control (Supplementary Table 2, Fig 2 b). The magnitude of R_1_ increase was proportional to the MAGIC MRI change, as expected due to the parameters of the MPRAGE sequence used. The largest R_1_ increases were observed in the ipsilateral GP, home to neuronal cell bodies sending their projections to the injected CP. These results indicated substantial increases in endogenous paramagnetic metals into Zip14-expressing areas. Of the metals known to be transported by Zip14, Mn^2+^ has the highest longitudinal relaxivity (r_1_)^5^. Cu^2+^ has substantially lower r_1_ than Mn^2+ 6^. Fe^2+^ can be transported by Zip14 and then oxidized and stored within ferritin in Fe^3+^ valence state with much lower T_1_ relaxivity than Cu^2+ 7^. Zn^2+^ is not paramagnetic^6^. Using previous measurements of Mn^2+^ r_1_ in tissue^5^, the change in measured R_1_ predicts a 1.5 – 2.5 ppm or roughly doubling of basal level Mn^2+^ concentration. Cu^2+^ and Fe^2+/3+^ have such low r_1_, that concentrations would need to increase by approximately 14 ppm for Cu^2+^ (135% of basal levels) or 375 – 10,000 ppm (400% - 11,500% of basal levels) for Fe^2+/3+^ to be responsible for the R_1_ change measured^7–9^. These estimates assume that the Fe^2+/3+^ is sequestered in ferritin and that the Cu^2+^ relaxivity is like that found in water.

LA-ICP-TOF-MS was applied to determine the quantitative change in metal ions.. It has been demonstrated that Zip14 transports Zn^2+^ and Cu^2+^ into cells in vitro, in addition to transporting Mn^2+^ and non-transferrin bound Fe^2+ 10,11^. However, in RetroAAV-Zip14 injected mice, no significant increases in Zn^2+^ concentration in the MAGIC hyperintense subregions of the CP nor in the ipsilateral GP nor SN were observed (Supplementary Figure 3). In the injected CP, Mn^2+^ concentration was found to be significantly increased and approximately double that of the contralateral control concentration (Fig 2 c, Supplementary Table 3). This 1.4 ppm increase, 83% increase from basal level, in concentration was in line with the predicted values from the observed change in R_1_ in the CP. The area of increased Mn detected in ex vivo LA-ICP-TOF-MS closely matched in vivo MAGIC hyperintensity and R_1_ increases, both in terms of spatial localization and geometry. To our knowledge, this is the first report of virally expressed Zip14 manipulating endogenous metal ion distribution in the brain in vivo.

**Fig 3.**
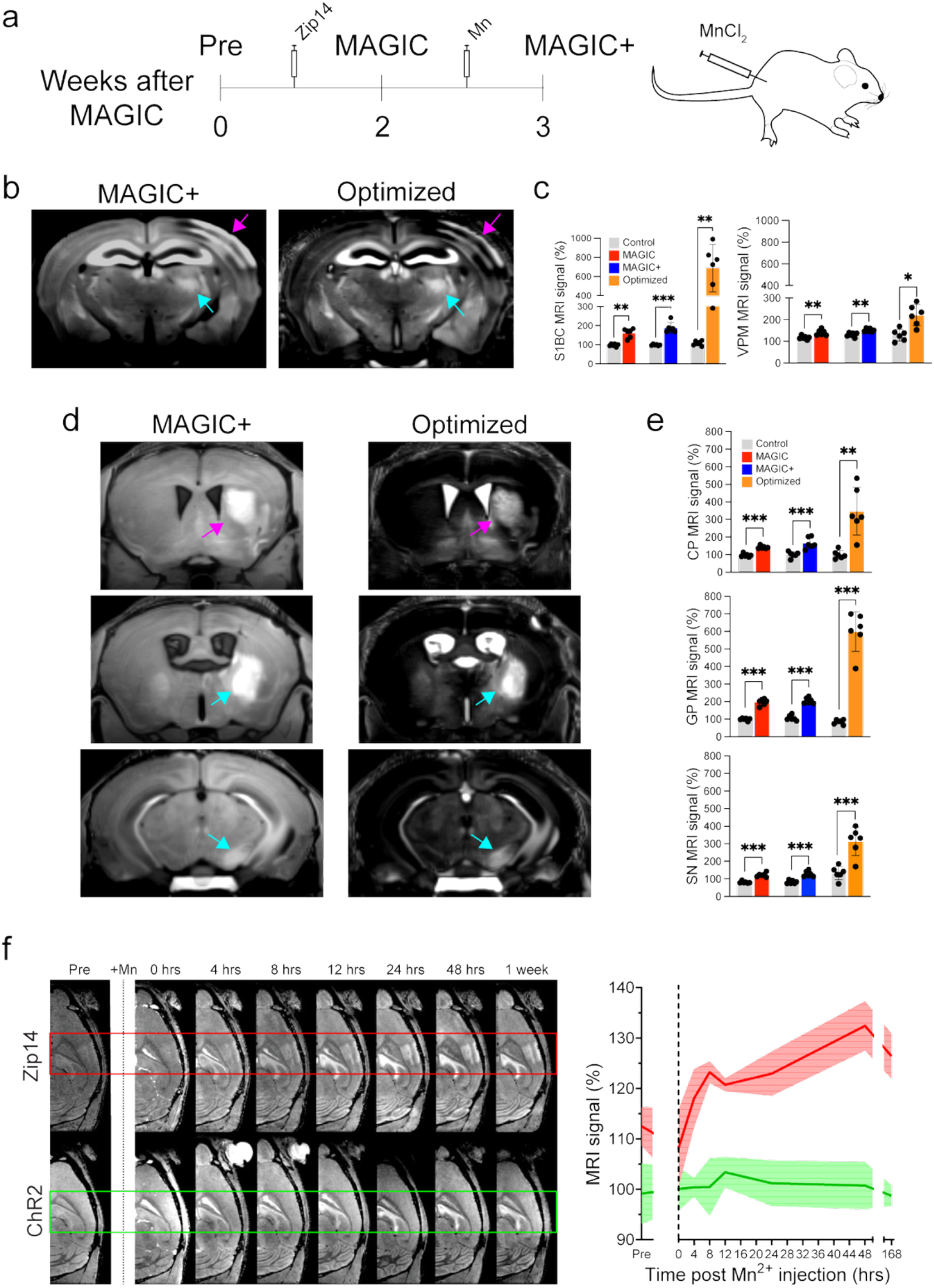
Mn^2+^ supplementation improves MAGIC (MAGIC+). (a) Animals previously injected with AAV-Zip14 or RetroAAV-Zip14 were intraperitoneally injected with MnCl_2_ one week after the MAGIC imaging timepoint. One day after supplementation, MAGIC+ was performed using the same MPRAGE protocol and an Optimized protocol with a shorter inversion delay to account for global T_1_ changes resulting from nonspecific Mn^2+^ uptake. (b) MAGIC+ images from an animal injected with AAV-Zip14 into the S1BC showed substantial signal intensity increase in MAGIC in the S1BC (magenta arrow) and ipsilateral VPM (cyan arrow). Quantitative contrast measurements were made, showing that these effects were reproducible across animals (n=6), robust, and statistically significant. (c) Optimization of the MPRAGE parameters improved contrast up to 5-fold for MAGIC+ in the injected S1BC. (d) MAGIC+ of animals injected with RetroAAV-Zip14 into the CP (magenta arrow) also displayed robust, statistically significant signal enhancement in the CP, GP (cyan arrow), and SN (cyan arrow) compared to conventional MAGIC. (e) The Optimized imaging protocol further improved MAGIC+, with the Zip14-expressing GP at 5-fold higher contrast than the contralateral control. (e) To test the persistence of the MAGIC improvement afforded by Mn^2+^ supplementation, timecourse images were acquired from animals injected with AAV-Zip14 (n=3) or AAV-Channelrhodopsin (n=3, ChR2) into the S1BC. MAGIC+ was significantly improved between Zip14 expressing areas and normal cortex compared to ChR2 for at least one week after supplementation (shaded area = 95% confidence interval). P-value thresholds: ^*^p<0.05 ^* *^p<0.01 ^* * *^p<0.001.

No significant increase in Cu^2+^ was measured in any MAGIC hyperintense brain area. In fact, an 18 ppm decrease in Cu^2+^ concentration, 54% of basal levels, was observed in the MAGIC bright subareas of the CP on average. Therefore, Cu^2+^ accumulation could not be the mechanism for MAGIC (Supplementary Figure 3).

LA-ICP-TOF-MS of snap frozen tissue proved to be well-suited for obtaining metal concentration measurements with spatial resolution similar to that of quantitative MRI maps, making it possible to estimate the in vivo r_1_ of Mn^2+^. 450 ROIs (320 µm diameter circles, 71 within MAGIC enhanced areas, 379 in normal appearing areas) across 3 animals were analyzed. These matched ^55^Mn metal maps and R_1_ mapping data revealed a strong linear correlation between Mn^2+^ concentration and R_1_ (Fig 2 d). The ROIs with highest R_1_ and Mn^2+^ were the high intensity MAGIC areas of the mouse brain in the Retro-AAV injected animals. Regression analysis gave an r_1_ value of 3.88 sec^-1^mM^-1^, similar to the range of 4 – 5 sec^-1^mM^-1^ for Mn^2+^ r_1_ in the brain reported previously^5,12^. A statistically significant 44 ppm increase in Fe^2+/3+^ concentration was measured, 150% of basal levels, in MAGIC bright subareas of the CP. (Fig 2 e, Supplementary Table 5,). Smaller magnitude increases in Fe^2+/3+^ were observed in the ipsilateral GP and SN. The largest magnitude Fe^2+/3+^ increases within the CP were localized to a smaller area than the increased Mn^2+^ and MAGIC hyperintensity; this increase was likely due to increased iron from bleeding at the injection site. However, and regardless of the source, the change in Fe^2+/3+^ concentration was several orders of magnitude (∼10 – 200x) too small to explain the observed change in R_1_, or the signal changes observed with T_1_ weighted MRI.

Given that Fe^2+/3+^ has poor longitudinal relaxation properties, R_2_^*^mapping was performed to determine if there was any potential for T_2_-weighted MAGIC caused by accumulation of Fe^2+/3+^ (Supplementary Figure 3). Fe^2+^ can be transported by Zip14 then be oxidized to Fe^3+^ and loaded into ferritin to exhibit the highest potential effective transverse relaxivity (r_2_^*^) of metals transported by Zip14. Voxel-wise correlation was performed between in vivo R_2_^*^data and matched ^56^Fe and ^55^Mn metal maps. Even in the extreme case where the entirety of the measured ^56^Fe was loaded into ferritin, the concentration change was too small to explain the very small R_2_^*^change in MAGIC enhanced areas. The concentration change in Mn^2+^ was a better predictor for the change in R_2_^*^based on published r_2_ values^5^. Therefore, these data indicate that the very small change in R_2_^*^is most likely due to Mn^2+^ concentration change.

### Mn^2+^ supplementation greatly improves MAGIC

Mn^2+^-enhanced MRI (MEMRI) has been useful to study neuroarchitecture, connectivity, and activity in the living rodent brain^13^. Increasing Mn^2+^ availability in the brain using systemic administration protocols routinely used in MEMRI studies has the potential to improve MAGIC (MAGIC+). Because MAGIC is highly correlated with Mn^2+^ concentration changes, supplementation would further increase confidence that increases in Mn^2+^ are responsible for the detected contrast. Animals injected with either AAV-Zip14 and RetroAAV-Zip14 into the brain, as in Fig 1, were injected intraperitoneally with 100 mM MnCl_2_ (64 mg/kg, 90 µL for a 30 g mouse) one week after the MAGIC scan session (Fig 3 a). Twenty-four hours after MnCl_2_ injection, animals were scanned using the MPRAGE protocol used for MAGIC and a modified MPRAGE protocol (Optimized) tailored to suppress gray matter signal throughout the brain after Mn^2+^ uptake.

Animals injected with AAV-Zip14 into the S1BC and intraperitoneally injected with MnCl_2_ exhibited substantially increased signal throughout the brain, consistent with the known widespread uptake of systemic Mn^2+^ into the brain^13–15^. Compared to the Mn^2+^ enhanced contralateral S1BC, the Zip14-expressing S1BC was reproducibly ∼90% higher signal level (Fig 3 b, Supplementary Table 6). Furthermore, the VPM ipsilateral to the injected S1BC was also enhanced by approximately 20% compared to the non-Zip14 expressing contralateral VPM. MAGIC+ at this projecting site was similarly robust and reproducible across animals (Supplementary Table 6). The optimized MPRAGE parameters to account for whole brain T_1_ shortening caused by global Mn^2+^ uptake markedly improved contrast at both the S1BC injection site and the projecting VPM (Supplementary Table 6, Fig 3 c). With optimized MAGIC+, the Zip14-expressing S1BC was approximately five times the signal level of the contralateral S1BC on average. The ipsilateral VPM had approximately double the signal level of the contralateral VPM on average (Supplementary Table 6).

MAGIC+ in RetroAAV-Zip14 injected animals improved contrast at the injected CP and basal ganglia subregions projecting to the ipsilateral CP. Zip14-expressing areas displayed MAGIC+ enhancement approximately equal to the effect of MAGIC (Fig 3 d,e). The injected CP and ipsilateral SN and GP were all significantly hyperintense, with the GP doubled in signal level compared to contralateral control areas. This effect was reproducible across animals, and the contrast improvements were statistically significant (Supplementary Table 6). The optimized MAGIC+ parameters substantially improved MAGIC+ across all Zip14 expressing areas. The effect of optimized MAGIC+ in the CP was nearly three times the signal level of the contralateral control. Optimized MAGIC+ in the GP was five times the signal level of the contralateral control. Optimized MAGIC+ in the SN was two times the signal level of the contralateral control (Fig 3 d,e, Supplementary Table 6).

To test the rate of increase of contrast due to Zip14 expression and systemic MnCl_2_ injection, time course imaging experiments were performed immediately after MnCl_2_ injection (Fig 3 d) for up to 1 week. Animals were injected with either AAV-Zip14 or AAV expressing Channelrhodopsin (ChR2) into the right hemisphere S1BC. Two weeks after virus injection and before MnCl_2_ injection, animals were scanned using an off-the-shelf gradient echo imaging protocol. This protocol was optimized to minimize acquisition time while maintaining high resolution, so some contrast-to-noise was sacrificed. Increased signal was observed in the injected S1BC of animals expressing Zip14 but not ChR2 as previously described^2^. This confirmed that MAGIC is produced by Zip14 expression and not due to non-specific effects of expressing membrane protein at high levels^2^. The ChR2 expressing S1BC and normal cortex remained consistently isointense across all imaging timepoints up to one week after MnCl_2_ injection. Contrast between the Zip14 injected S1BC and surrounding normal appearing gray matter was slightly (∼10%) reduced immediately after MnCl_2_ injection, likely due to increasing Mn^2+^ concentration in the cerebrospinal fluid^14^. Signal was significantly improved over Pre-baseline four hours after MnCl_2_ injection and continued to increase up to 48 hours post injection where it peaked at roughly double the contrast enhancement prior to injection. Zip14 expressing areas remained hyperintense above Pre-baseline and ChR2 expressing areas one week after MnCl_2_ injection.

### Fully automated detection of MAGIC

Exploiting the marked improvement in MAGIC afforded by MAGIC+, an automated enhancement detection pipeline was developed using anomaly mapping of an individual brain compared to an average of control brains (Fig 4). This procedure enables analysis of a single subject given a large enough group size of average brains, analogous to analysis pipelines processing UKBiobank and the Human Connectome Project data^16,17^, assuming relatively large effect size on the individual difference. Template images from a relatively large population of naïve mice were constructed. Mice were either completely unperturbed (n = 24) or intraperitoneally injected with MnCl_2_ (n = 20) and scanned using the off-the-shelf MPRAGE sequence used for conventional MAGIC experiments. It should be noted that for these experiments the optimized MPRAGE imaging protocol was not used in template image construction nor anomaly detection. The heavily suppressed normal-appearing gray matter signal in the optimized MAGIC+ images resulted in poor intersubject brain registration. Images were nonlinearly registered together and subsequently averaged to produce MRI and MEMRI template images. To detect anomalous signal compared to the control template in MAGIC or MAGIC+ images obtained from an AAV-Zip14 or RetroAAV-Zip14 injected mouse, images were nonlinearly registered to the appropriate template space, the registered image signal was globally shifted such that the mean matched the template mean, and voxel-wise z-scoring was performed (Fig 4 a). Before testing MAGIC or MAGIC+ images, leave-one-out cross validation was performed on the naïve data used to produce each template to determine the false positive detection rate at increasing z-score thresholds (Fig 4 b). For z-scores equal to two standard deviations above the mean or greater, the MAGIC+ pipeline was significantly less prone to false positives than MAGIC alone. Both MAGIC+ and MAGIC pipelines had false positive detection rates less than 5% for z-scores equal to or greater than three, and false positive detection rates less than 1% for z-scores equal to or greater than four. Whole-brain anomaly detection was performed for RetroAAV-Zip14 injected animals using both MAGIC and MAGIC+ data. A representative RetroAAV-Zip14 injected animal’s MAGIC anomaly map revealed high z-scores in the injected CP and ipsilateral GP and SN (Fig 4 c). The highest z-scores were detected in the GP. Other areas with moderately high z-score, but sparse spatial distribution included the colliculus and medial-dorsal cortical areas. The MAGIC+ anomaly map from the same animal showed higher z-scores in the injected CP and areas known to project to it (Fig 4 d). However, the spuriously detected regions observable using the MAGIC anomaly detection pipeline were not detected in the MAGIC+ anomaly map: further evidence for increased specificity afforded by MAGIC+. MAGIC+ anomaly maps were visually compared to mesoscale connectivity data available in the Allen Mouse Brain Connectivity Atlas^18^ with good agreement.

**Fig 4.**
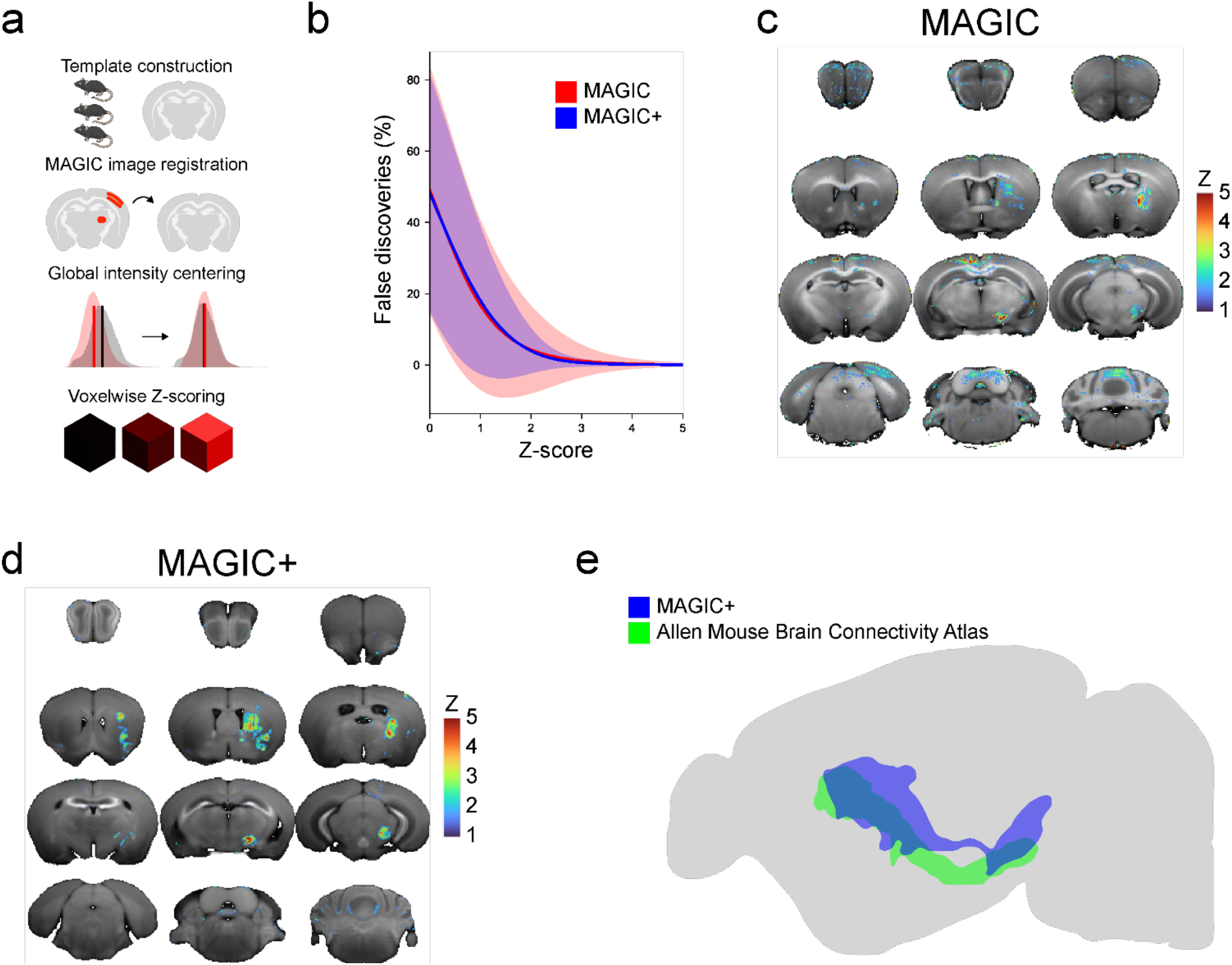
Automated whole-brain MAGIC detection. Schematic of the image processing pipeline. First, large sample size cohorts of naïve and Mn^2+^-supplemented mice were imaged with the conventional MPRAGE protocol used for MAGIC. These images were registered together to create template images and standard deviation maps. Next, MAGIC and MAGIC+ images were registered into their respective template space and intensities were linearly centered to match mean values with the template. Finally, voxelwise z-scoring was performed to test if MAGIC and MAGIC+ voxels were significantly different from normal. (b) Leave-one-out cross validation of the image processing pipeline for naïve and naïve MEMRI data. Compared to conventional MAGIC data, the pipeline applied to MAGIC+ data produced significantly fewer false positive high z-score detected voxels. (c) Representative whole brain enhancement map for MAGIC data acquired from a RetroAAV-Zip14 injected animal shows high z-score values in the CP, GP, and SN. Spurious detections in the medial anterior cortex and cerebellum were also observed. (d) The improved signal-to-noise afforded by MAGIC+ of the same animal reduced the false-positive detections while also increasing the effect size in the Zip14-expressing areas. (e) Comparison of whole brain detected MAGIC+ to mesoscale connectivity data matched by injection site from the Allen Mouse Brain Connectivity Atlas shows close spatial colocalization.

### MAGIC in rhesus macaques

Many MRI visible gene expression reporter systems have been developed, and several have shown promise for neuroimaging applications^1^. However, none have yet crossed the threshold for clinical translation nor have demonstrated efficacy in non-human primates. To determine if MAGIC or MAGIC+ was sufficiently robust for use in non-human primate studies, they were tested in rhesus macaques (Fig 5). A spectrum of capsids, viral titers, injection sites, and animal ages were tested. Lentivirus, AAV2, AAV9, and RetroAAV were packaged with plasmid constructs to express Zip14 under the control of the Synapsin promoter. Stereotaxic injections were performed targeting the striatum (caudate or putamen) and the cortex. All animals were imaged once before Zip14 viral injection, and serially up to four months after injection. Then all animals, except the one injected with RetroAAV-Zip14 into the striatum, were supplemented with MnCl_2_ delivered through the saphenous vein. Follow up MAGIC+ was performed weekly up to two weeks after systemic MnCl_2_ administration (250 mM, 10 mg/kg, 2 mL for a 10 kg macaque) to the animals (Fig 5 a). AAV2-Zip14 injections did not produce any in vivo MAGIC or MAGIC+ enhancement nor positive Zip14 staining in matched immunofluorescence histology across either injection site. Similarly, none of the injections into the cortex generated positive immunohistology staining nor MAGIC or MAGIC+. RetroAAV-Zip14 injections produced marked MAGIC enhancement and Lentivirus and AAV9-Zip14 injections into the striatum caused robust MAGIC+ enhancement in vivo. These in vivo imaging findings were validated with positive Zip14 staining in immunofluorescence histology. It is not clear why some capsids worked and some did not, nor why there was no enhancement from the cortical injections.

**Fig 5.**
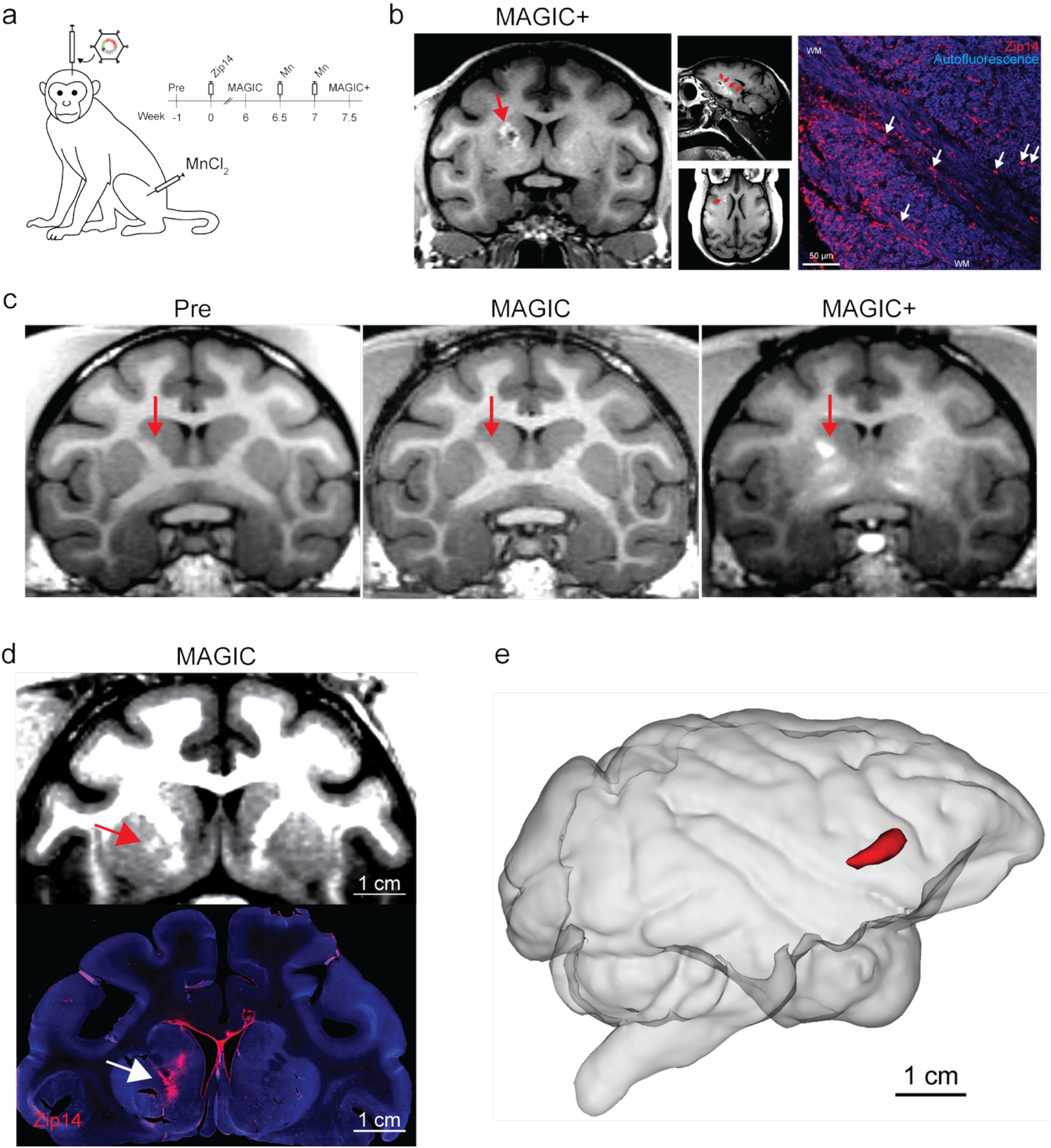
MAGIC in rhesus macaques. (a) Schematic non-human primate MAGIC methods. 2 male and 2 female macaques were injected with combinations of AAV2, Lentivirus (Lenti), RetroAAV, and AAV9 expressing Zip14 under the control of the Synapsin promoter. Stereotaxic injections targeted the caudate and putamen and cortical sites, however none of the cortical injections produced any Zip14 expression. Of these constructs, Lenti-Zip14 and RetroAAV9-Zip14 were most performant; they produced MAGIC enhancement and expression confirmed by matched immunohistology. (b) Three-axis views of 500 µm isotropic resolution MEMRI from a macaque injected with Lenti-Zip14 showed bright signal in the caudate, putamen, and internal capsule around a small edematous lesion (red arrow). This enhancement was colocalized with high Zip14 expression in neuronal cell bodies, dendrites, and axons (white arrows) in the striatal gray matter and white matter (dashed lines, WM) produced by the delivered lentivirus. (c) Representative Pre scans, MAGIC scans, and MAGIC+ images from another animal injected with Lenti-Zip14 into the caudate and AAV9 into the putamen. No enhancement was apparent on Pre and MAGIC images, but MAGIC+ focal enhancement centered on the internal capsule and extending into the caudate and putamen. (d) Surprisingly, RetroAAV-Zip14 injections produced detectable MAGIC (red arrow) in another macaque without requiring supplemental Mn^2+^. This enhancement was also colocalized with Zip14 expression produced by the viral vector. Because of the similarity in contrast between the normal appearing gray matter and MAGIC enhancement observed in this animal, the MRI image is displayed with in a compressed intensity window compared to the representative MAGIC+ images. (e) The high contrast afforded by MAGIC+, as shown in (b), enabled 3D rendering of enhancement in the macaque brain.

Three-axis views through a MAGIC+ image acquired from a monkey injected with Lentivirus-Zip14 into the left hemisphere striatum and AAV2-Zip14 into the right hemisphere showed bright, focal enhancement only in the left hemisphere (Fig 5 b). Sagittal and axial views at the level of the injection site also revealed that the basal ganglia in the left hemisphere (ipsilateral to the lentivirus injection site) were MAGIC+ hyperintense. Immunofluorescence histology for Zip14 confirmed that the enhancement was localized to Zip14 expressing areas driven by viral transduction of neurons in the left hemisphere striatum. Representative images from before and one month after injection of AAV9- and Lentivirus-Zip14 into the striatum (Pre, MAGIC) in one male monkey did not show any noticeable MAGIC enhancement at the injection site (red arrow, Fig 5 c). Bright, focal enhancement was apparent in the striatum on the MAGIC+ image acquired two weeks after the MAGIC scan (Fig 5 c). Focal MAGIC enhancement was apparent in a RetroAAV-Zip14 injected animal without supplemental MnCl_2_ administration (Fig 5 d). Matched immunofluorescence histology confirmed MAGIC enhancement was localized only to Zip14 expressing areas near the RetroAAV injection site in the striatum. The strong and reproducible signal enhancement afforded by Zip14 expression driven by viral vectors enabled 3D rendering of in vivo MAGIC in the macaque brain (Fig 5 e).

## Discussion

This study shows that MAGIC is a versatile GEMRI platform technology for neuroscience. Consistent with results obtained in previous studies, application of the Mn^2+^ transporter Zip14 expressed using viral vectors enabled non-invasive, mesoscale neural tracing in vivo in the rodent^2^. The present work demonstrates that the mechanism for MAGIC is redistributing endogenous Mn^2+^ towards Zip14 expressing areas. Understanding the contrast mechanism enabled MAGIC to be amplified with supplemental Mn^2+^ (MAGIC+). An image processing and analysis pipeline was developed to automatically detect and quantify MAGIC and MAGIC+ enhancement in individual animals. Furthermore, the translational potential of MAGIC was established by demonstrating its efficacy in rhesus macaques, a first for any GEMRI system.

MAGIC is sufficiently robust for mapping neural circuits in the rodent brain in vivo^2^. By packaging Zip14 with different viral capsids, both anterograde projections (from S1BC to VPM with AAV9) and retrograde connections (from GP and SN to the CP with RetroAAV) were successfully visualized. This flexibility allows for the targeted enhancement and tracing of specific pathways in vivo. For S1BC injections, heterogeneity of this bright signal across the cortical depth indicated cortical layer-specificity of Zip14 expression, a product of the well-established neurotropism of AAV9 in mice^19^. In future studies, it will be interesting to determine whether the MRI contrast is related in a quantitative way to numbers of cells expressing Zip14 and/or to axonal arbor density especially in anterograde sites. That Zip14 expression was the source of MAGIC was confirmed through immunofluorescence histology, which verified that the hyperintense signals on MRI were specifically colocalized with high levels of Zip14 expression in neurons. This validation provides confidence that MAGIC is a reliable reporter of gene expression and, when expressed in neurons, anatomical connectivity. If the Mn^2+^ loaded by Zip14 expressing neurons is able to cross the synapse as is the case with direct injections of MnCl_2_ into the brain^12^ (i.e., is not cell autonomous), then there is potential for MAGIC detecting connectivity across multiple synapses.

A central finding of this study is the association of MAGIC to increased Mn^2+^ concentration. In situ detection of Mn^2+^ using ex vivo assays has been challenging because redistribution occurs postmortem, and no histological stain currently exists^20^. Through quantitative R_1_ and R_2_^*^ mapping and high-resolution LA-ICP-TOF-MS, it was determined that Zip14 produces MAGIC consistent with the increase in total Mn^2+^. The strong linear correlation between local Mn^2+^ concentration and R_1_ values makes it likely that Zip14 concentrated Mn^2+^ to levels sufficient to generate robust T_1_-weighted contrast without any external agents. While Zip14 also transported Fe^2+^, the change in Fe^2+/3+^ concentration was too small to contribute to the overall T_1_ weighted MAGIC MRI signal change. It is likely that needle-induced local bleeding from the stereotaxic injection could have resulted in much of the observed increase in Fe^2+/3+^ in the injected CP since no significant changes in iron occurred in GP or SN. In short, Mn^2+^ was the only metal to increase in concentration in all three areas (CP, GP, and SN) and no other metal concentrations changed sufficiently to create the observed MRI contrast change.

Zip14 has been shown to efficiently transport metal ions other than Fe^2+^ and Mn^2+^, including Cu^2+^ and Zn^2+^, in vitro^21^. Though these experiments exposed cultured cells to near-physiological concentrations of these divalent metals, our in vivo results showed no significant increase in Zn^2+^ and a modest drop in Cu^2+^ in Zip14 expressing areas. Furthermore, in vitro metal competition assays performed using Zip14-expressing cells revealed that exposure of Mn^2+^ inhibited transport of many divalent metals^21^ establishing Mn^2+^ as the metal with highest affinity for Zip14 in vivo when expressed in neurons. It is possible that that compensatory transport of other metals may occur, however the fact that concentrations of other metals did not change substantially indicates that the effect on metal homeostasis is not large. Human and mouse mutations of Zip14 demonstrate that it is a critical regulator of systemic Mn^2+^. Loss of function mutations in Zip14 block the uptake of Mn^2+^ by the liver, preventing its excretion into bile^3,4,22^. This results in severe hypermanganesemia and subsequent toxic accumulation of Mn^2+^ in the brain^3^. Taken together, these results indicate that Mn^2+^ accumulation mediated by Zip14 transport the principal driver of MAGIC.

While MAGIC is effective using only endogenous Mn^2+^, its signal can be reliably increased by systemic administration of MnCl_2_, a common agent for MRI used in preclinical neuroimaging studies. Significantly improved sensitivity to Zip14 expression was afforded by MAGIC+, including increasing contrast by up to five fold in the GP of RetroAAV-Zip14 injected mice when using optimized MPRAGE parameters. This enhancement was shown to be rapid, peaking at 48 hours and persisting for at least one week, providing a flexible and extended window for imaging. Mn^2+^ dosage of 64 mg/kg was chosen to match previous MEMRI studies^23–25^. We have tested doses as low as 30 mg/kg and obtained proportional MAGIC+ enhancement, indicating lower doses of MnCl_2_ may be sufficient (Supplementary Figure 4). In this study, no attempt was made to determine the rate limiting steps to contrast generation. Nevertheless, the fact that supplementation with systemic MnCl_2_ increased contrast indicates that Mn^2+^ transport into Zip14 expressing areas may be one of the limiting factors.

This potent contrast boost enabled the development of a fully automated MAGIC detection pipeline. The high contrast-to-noise ratio produced by MAGIC+ allowed the developed algorithm to identify Zip14-expressing regions with a false-positive rate of less than 1% (at z > 4). The high z score required to make decisions on an individual compared to a group average indicate that retrograde strategies to express Zip14 are likely to resolve mesoscale connectivity in a single subject. The variance in false positive detected voxels using the MAGIC+ data were significantly reduced compared to conventional MAGIC data. Automated anomaly detection will be useful for future high-throughput screening of circuit alterations in disease models or therapeutic interventions, and/or for initiating MAGIC MRI studies at new imaging sites. The current pipeline relies on image registration and image subtraction; a relatively simple method that functions because of the large effect size afforded by MAGIC and MAGIC+. The well-curated dataset of naïve control and MAGIC images produced through this work is in shape to train deep learning models to score subtle MAGIC MRI changes, such as enhancement from sparse Zip14 expressing presynaptic terminals^16,26–28^.

The successful demonstration of MAGIC in the rhesus macaque at clinical MRI field strengths lays the foundation for translational application of MAGIC, and should make in vivo, mesoscale gene expression studies in nonhuman primates widely accessible. Presently, most in vivo gene expression analyses in macaques have required the use of PET^29^. The expense and lack of accessibility to the required specialized radiolabeled agents have hampered widespread application. Furthermore, the low spatial resolution afforded by PET compared to the relatively small size of the non-human primate brain may have hindered more studies. MAGIC MRI is a strong candidate alternative to PET expression reporter strategies because it potentiates multiple contrasts, nonionizing radiation in acquisition, without radioisotopes, sensitivity unlimited by tissue depth, and relatively high resolution. In this study, it was found that Lentivirus, AAV9, and RetroAAV induced robust, detectable MAGIC in the macaque striatum. It is not clear why cortical injections of the virus did not produce expression of Zip14 and thus no MAGIC. Technical problems with injecting into the thin cortex could be the cause, though there are many reasons why MAGIC is more apparent in white matter-dense areas. The first is that the cell bodies expressing Zip14 are relatively sparse compared to their axons which concentrate to relatively high density in the white matter. Therefore, there may be a more uniform increase in Mn^2+^ in the white matter compared to the gray matter. Next, the MPRAGE protocol used is relatively low resolution compared to the cell body diameter, density, and Zip14 expressing fraction. It is likely that partial volume-effects drive a decrease in contrast in the gray matter. As this protocol is optimized for gray matter/white matter contrast in a relatively short acquisition time, some parameters lead to “blurring” of tissues with different magnetization relaxation properties. Nevertheless, the fact that in most cases MAGIC+ MRI had to be performed gives is evidence that transport of Mn^2+^ to Zip14 expressing areas is a rate limiting step to contrast in the macaque brain. Differential Mn^2+^ uptake after systemic supplementation (e.g., high uptake in periventricular areas, as has been shown in marmosets) may also affect detection of Zip14 expression in low-availability brain areas^30^. However, the ability to detect neuronal populations in a living primate brain using a GEMRI system, even without supplemental Mn^2+^ in one case, is unprecedented. This result establishes MAGIC as a translationally relevant platform technology and opens the door for studying large-scale circuit dynamics in higher-order animals whose brain organization more closely resembles that of humans. In summary, this study establishes MAGIC as a robust, scalable, and translationally relevant platform for mapping neural circuits in living brains across species.

## Methods

### Animals

All rodent procedures were approved by the NINDS Animal Care and Use Committee under protocol 1142, and facilities are accredited by the Association for Assessment and Accreditation of Laboratory Animal Care. C57Bl/6J mice were bred in house, originally purchased from the Jackson Laboratory (Strain ID: 000664). Animals of both sexes were used, and no significant differences in imaging results were detected between sexes. Mice were housed up to four per cage with food and water ad libitum on a 12/12 h light/dark cycle.

Animals were 6–8 weeks old at the time of surgery. They were anesthetized with 1%–3% isoflurane mixed with 40% O_2_ in compressed air, a hole was drilled in the skull, and then unilaterally injected at a rate of 100 nL/min using a pump (World Precision Instruments;UMP3T-1) and syringe (Hamilton #1701 body, #7803-07 30 GA 45◦ needle) into various brain areas including the primary somatosensory cortex, barrel field (S1BC), ventral posteromedial nucleus of the thalamus (VPM), and the caudate putamen (CP).

Macaque experimental procedures followed the Institute of Laboratory Animal Research Guide for the Care and Use of Laboratory Animals and were approved by the National Institute of Mental Health Animal Care and Use Committee. The virus injection was performed on four monkeys: Monkey P (weight: 11.1 kg, age: 9 yrs, sex: male), Monkey D (weight: 10.3 kg, age: 19 yrs, sex: female), Monkey G (weight: 8.8 kg, age: 7 yrs, sex: male), and Monkey Z (weight: 7.3 kg, age: 13 yrs, sex: female). Before the operation, the animal was placed in a stereotaxic frame and a 3.0 T MR image was obtained to determine the cortical and striatal injection sites. The monkey was then transported to the operating room in the stereotaxic frame. Surgical procedures were performed using sterile technique under general anesthesia in a fully equipped operating room with veterinary supervision. To allow the injection apparatus to access the injection sites, the skin, fascia, and temporal muscle were reflected. A bone flap was removed, and a small incision was made in the dura mater underneath. The virus was loaded into a 100 µL glass syringe (Hamilton Co., MA) that was affixed to a nanomite pump (Harvard Apparatus, Cambridge, MA). The syringe was lowered to the target sites. The infusion rate in cortical regions was 0.5 µL/min and 1 µL/min for striatal regions. After infusion, the needle was left for 10 minutes in the injection site before being slowly lifted. For Monkey P, 60 µL of retro-AAV9 with a titer of 6×10^12^ vg/mL was injected into the rostral medial caudate of the left hemisphere. For Monkey D, Lentivirus and AAV2 at a titer of 4×10^9^ IU/mL and 10^12^ vg/mL respectively, were used. Lentivirus was injected in the left hemisphere (40 µL in the caudate and 20 µL in the cortex). AAV2 was injected in the right hemisphere (40 µL in the putamen and 20 µL in the cortex). For Monkey G and Monkey Z, 40 µL of lentivirus at a titer of 2×10^9^ IU/mL was injected into the caudate and 20 µL into the cortex of the left hemisphere. AAV9 was injected in the putamen at a volume of 35 µL, and 15 µL was injected in the cortex of the left hemisphere at a titer of 6×10^12^ vg/mL for Monkey G and 6×10^13^ vg/mL for Monkey Z.

### MRI

Small animal MRI were performed before, immediately after, 2 weeks after, and many timepoints after injection of AAV-Zip14 using a Bruker 11.7 T MRI system, Bruker Avance NEO system, RRI or Bruker gradient systems, and a Cryoprobe. A conventional T1 weighted, 80 µm isotropic resolution MPRAGE protocol was used to produce most of the presented data. Acquisition parameters: echo time, 3.6 ms; repetition time, 4000 ms; segments, 2; flip angle, 12°; inversion time, 1100 ms; bandwidth, 50 kHz; matrix size, 176 × 176 × 176; FOV, 14 × 14 × 14 mm; averages, 2; and acquisition time, 47 minutes. The exception was the Mn^2+^-supplemented imaging timecourse, for which a standard T1 weighted, 80 µm isotropic resolution FLASH sequence was used. Acquisition parameters: echo time, 4.6 ms; repetition time, 30 ms; flip angle, 20°; bandwidth, 50 kHz; matrix size, 196 × 196 × 96; FOV, 15.68 × 15.68 × 7.68 mm; averages, 2; and acquisition time, 18 minutes.

Three weeks after AAV-Zip14 injection, 100 mM MnCl_2_•4H_2_O was delivered intraperitoneally at least 20 hours before MAGIC+ experiments (60 – 100 µL, 64 mg/kg) as previously described^31^.

Macaque MRI were performed before, immediately after, 1 week after, and many timepoints up to 4 months after the injection of Zip14-expressing viral vectors using a Siemens 3 T MRI system, Prisma Syngo E11C system, Siemens XR gradient, and a 11 × 10 cm oval custom-made single loop coil. All the present data were acquired in the headfirst prone position starting with a 250 × 250 mm field of view localizer followed by a 0.5 × 0.5 × 0.5 mm isotropic resolution T1 weighted MPRAGE. Volumes of 224 slices were collected in the sagittal orientation, with a field of view of 112 × 112 mm on a 224 × 224 matrix with a slice thickness of 0.5mm. The acquisition parameters were echo time, 2.22 ms; repetition time, 2550 ms; turbo factor, 196; flip angle, 9°; inversion time, 900 ms; bandwidth, 290 Hz/pz; and acquisition time, 12 minutes.

For fractionated Mn^2+^ supplementation in rhesus macaques, MnCl_2_∙4H_2_O was administered at a concentration of 250 mM for a total dose of 10 mg/kg of via the saphenous vein at a rate of 0.5 mL/min. These intravenous injections were performed four months after Zip14-expressing viral vectors were injected for Monkey D and 7 weeks after viral injections for Monkey D and Monkey G at the same concentration and rate. For all monkeys, a second MnCl_2_∙4H_2_O injection was administered exactly 1 week after the first. MAGIC+ experiments took place 48 hours after the second supplementation using the same T1 weighted MPRAGE protocol.

### LA-ICP-TOF-MS

For LA-ICP-TOF-MS analysis of metal concentrations: animals were euthanized, brains extracted and snap frozen, cryosectioned at 20 µm thickness directly to glass slides, stored at -80 ºC until 24 hours before ablation where they were transferred to -20 ºC. Brightfield images were acquired of all slides using an Axioscan 7 slide scanner. A test brain section was run at 50, 60, 70, 75, and 80% laser power (1308/northwestern system) - 70% power was determined to be optimal. Analysis was performed using 20 µm circular spot size, 20 µm raster spacing, no overlap, at 100 Hz repetition rate using an ESL 266 Bioimage laser ablation system interfaced with a Tofwerk S2 ICP-TOF-MS at the Michigan State University QBEAM Facility.

### Viruses

Adeno-associated viruses were produced by the NINDS Viral Vector Production Core facility and by VectorBuilder. The Mn^2+^ transporter Zip14 was expressed using a plasmid construct consisting of a Synapsin promoter for neuronal specific expression, *Slc39a14* isoform 1, and a histology visible tag (either EGFP or FLAG) designed to be added to the C-terminus of Zip14. The plasmid was packaged into either AAV9 or RetroAAV capsids for anterograde (AAV-Zip14) or retrograde (RetroAAV-Zip14) labeling, respectively, with final working titers of up to 3×10^13^ vg/mL.

To create Lenti-Zip14, the hM4Di-CFP sequence was removed from pLenti-syn::hM4Di_CFP by digesting the plasmid with enzymes XbaI and EcoRV, and was replaced with the ZIP14-FLAG sequence from the pAAV hSyn1.SLC39A143xFLAG plasmid. In-Fusion cloning (638948, Clonetech) was used to combine the vector and fragment.

Lentivirus was produced by cotransfecting Lenti-X 293T cells with the Lenti-Zip14 plasmid, helper plasmid (psPAX2), pMD2.G, and transfection enhancer, P-AdVAntage. The supernatant was replaced 24 h later with new medium and Ultraculture (BP12-725F; Lonza). Lentivirus was collected 48 h after transfection by centrifuging supernatant in a 20% sucrose cushion in PBS at 22,000 rpm for 2 h. Finally, the pellet was suspended in PBS, aliquoted, and placed in -80 ºC for long-term storage. The virus was titered by thawing the aliquots on ice, then using 3µl of the virus at different dilutions (1:1, 1:10, and 1:100) to transduce 1.5 × 105 Lenti-X 293T cells. Genomic DNA was extracted from the cells 48 h later. Finally, qPCR was performed for validation.

### Histology

After the last imaging session, mice were deeply anesthetized with isoflurane and transcardially perfused with 10% formalin in phosphate-buffered saline using a peristaltic pump (Cole-Parmer, Vernon Hills, IL, USA). Brains were dissected and postfixed in 10% formalin overnight. Then they were immersed in 30% sucrose in phosphate-buffered saline until sunk and then snap-frozen in OCT in a dry ice and isopentane bath. 20 μm-thick coronal sections were cut using a cryostat (CM3050 S; Leica, Deer Park, IL, USA) and stored in Trizma-buffered saline and 0.1% sodium azide solution. Sections of interest were blocked with animal-free blocker (SP-5035; Vector Laboratories, Newark, CA, USA) for 1 h, then immunoreacted with primary antibodies at 1:200 volume dilution overnight at 4º C. The sections were then washed with Trizma-buffered saline and immunoreacted with secondary antibodies, as appropriate, at 1:200 volume dilution for 1 h at room temperature. Finally, sections were washed with Trizma-buffered saline, mounted on Apex Superior Adhesive Slides (#3800080 White; Leica), dried overnight at room temperature, and protected with ProLong Gold antifade reagent with DAPI (P36931; Invitrogen) using 24 × 50 mm #1 cover glass (EF15972LZ; Daigger).

Rhesus macaques were euthanized following AMVA guidelines and perfused with 1 L of normal saline solution, followed by a solution of 4% paraformaldehyde in 0.1 M phosphate buffer. The brains were removed and cryoprotected through a series of glycerol solutions in 0.1M PBS. The tissue was then frozen in isopentane and serially sectioned into 40 µm slices along the coronal plane using a sledge microtome. For histochemistry, sections were first placed in 4% paraformaldehyde for 30 minutes. The sections were then washed and blocked in 5% normal goat serum and 0.3% Triton-X 100 in PBS, incubated with primary antibody (HPA0165008; Sigma-Aldrich) at 1:200 volume dilution for 72 h at 4 ºC, washed, and incubated with secondary antibody (A21428; Invitrogen) at 1:200 volume dilution for 2 h at room temperature. Sections were washed a last time with PBS and mounted on 50 × 75 × 1.2 mm microscope slides (#5075, Brain Research Laboratories). Mounting media (18606-5, Polysciences) was added 20 minutes later, and slides were cover slipped using a 48 × 65 mm #1 cover glass (#4865-1D, Brain Research Laboratories).

### Statistics and Data Processing

Quantitative and semi-quantitative data analyses were performed using Python and GraphPad Prism. For region-of-interest (ROI)-based quantification of MAGIC and MAGIC+ changes, an ROI was manually drawn within the appropriate anatomical boundary based on Allen Institute Mouse Brain Atlas (CCFv3) definitions. ROIs drawn in the left hemisphere (non-injected, control) S1BC across all animals within an injected cohort was used to normalize signal measurements to 100%. For ROI-based quantification of MR parameter maps (R1, R2*), a similar method was employed. ROIs within the CP, GP, and SN were drawn within the anatomical boundaries for each region defined by the CCFv3 atlas.

To quantify metal concentration changes due to Zip14 expression in LA-ICP-TOF-MS maps, an automated ROI detection pipeline was developed. Briefly, raw metal map images were median filtered to remove streaking artifacts, the magnitude of the intensity gradient was calculated, a threshold was set to include only gradients above the 95^th^ percentile, the largest connected component was selected, and finally a convex hull was drawn around this component. The ROI was reflected to the contralateral hemisphere for basal metal concentration measurements.

Automated MAGIC and MAGIC+ detection consisted of three distinct phases: template construction, test image registration and normalization, and voxel-wise z-score calculation. MPRAGE images from naïve mice (n = 24) and MnCl_2_-injected mice (n = 20) were nonlinearly registered together using the SyN algorithm exposed in ANTs^32^ to create naïve and MEMRI average images and standard deviation maps. MAGIC or MAGIC+ images were nonlinearly registered to the appropriate average image. For each registered MAGIC or MAGIC+ image, the intensities of each voxel were shifted to match the template mean intensity. Then voxel-wise subtraction from the average template and division by the standard deviation map was performed to produce in voxel-wise z-score maps.

## Supporting information

LA-ICP-TOF-MS supplementary methods

## Acknowledgements

This work was supported by the Intramural Research program of the National Institutes of Neurological Disorders and Stroke (NINDS), National Institutes of Health. We thank Raymond Fields in the NINDS Viral Production Core Facility for virus production. Functional and anatomical MRI scanning of nonhuman primates was carried out in the Neurophysiology Imaging Facility Core (NIMH, NINDS, NEI).

## Supplementary Tables

**Supplementary Table 1.**
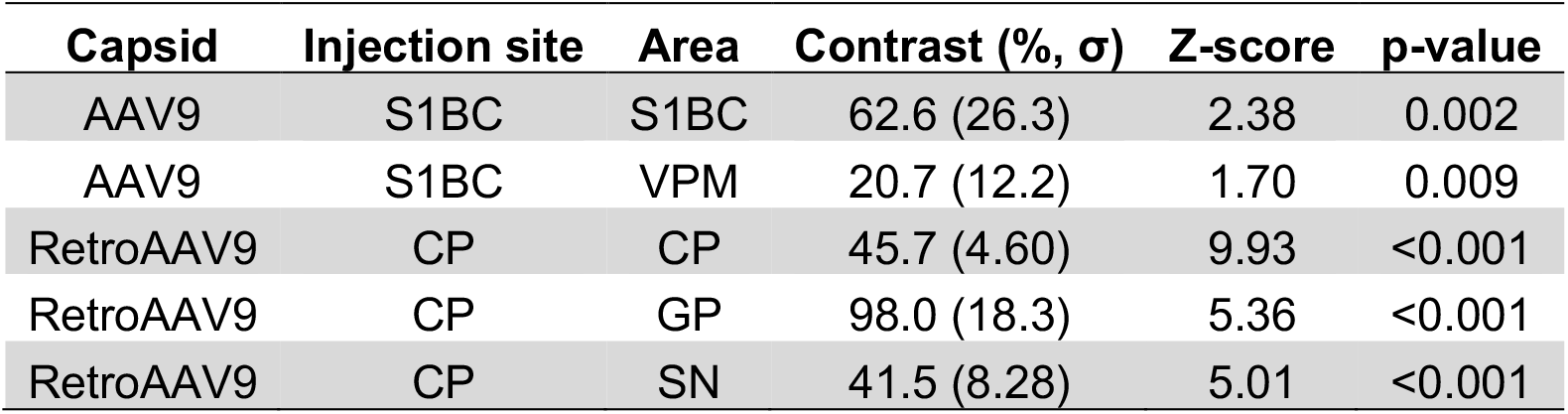
MAGIC effect size calculations.

| Capsid | Injection site | Area | Contrast (%$\sigma$) | Z-score | p-value |
| --- | --- | --- | --- | --- | --- |
| AAV9 | S1BC | S1BC | 62.6 (26.3) | 2.38 | 0.002 |
| AAV9 | S1BC | VPM | 20.7 (12.2) | 1.70 | 0.009 |
| RetroAAV9 | CP | CP | 45.7 (4.60) | 9.93 | <0.001 |
| RetroAAV9 | CP | GP | 98.0 (18.3) | 5.36 | <0.001 |
| RetroAAV9 | CP | SN | 41.5 (8.28) | 5.01 | <0.001 |

**Supplementary Table 2.** MAGIC quantitative R1 (sec^-1^) mapping results.

| Area | Control R1 ($\sigma$) | MAGIC R1 ($\sigma$) | ΔR1 | Z-score | p-value |
| --- | --- | --- | --- | --- | --- |
| CP | 0.691 (0.043) | 0.838 (0.030) | -0.147 (0.053) | -2.79 | 0.001 |
| GP | 0.741 (0.061) | 1.09 (0.080) | -0.352 (0.067) | -5.26 | <.001 |
| SN | 0.827 (0.071) | 1.02 (0.041) | -0.190 (0.098) | -1.94 | 0.005 |

**Supplementary Table 3.** MAGIC LA-ICP-TOF-MS ^55^Mn concentration (ppm)

| Area | Control [<sup>55</sup>Mn] ($\sigma$) | MAGIC [<sup>55</sup>Mn] ($\sigma$) | $\Delta$[<sup>55</sup>Mn] ($\sigma$) | Z-score | p-value |
| --- | --- | --- | --- | --- | --- |
| CP | 1.69 (0.203) | 3.10 (0.405) | -1.41 (0.210) | -6.71 | 0.007 |
| GP | 1.53 (0.406) | 2.75 (0.345) | -1.22 (0.223) | -5.47 | 0.011 |
| SN | 1.89 (0.579) | 3.52 (0.432) | -1.63 (0.345) | -4.72 | 0.014 |

**Supplementary Table 4.** MAGIC quantitative R2* (sec^-1^) mapping results.

| Area | Control R2* ($\sigma$) | MAGIC R2* ($\sigma$) | ΔR2* ($\sigma$) | Z-score | p-value |
| --- | --- | --- | --- | --- | --- |
| CP | 38.0 (2.05) | 38.8 (3.03) | -0.793 (2.15) | -0.369 | 0.407 |
| GP | 42.2 (5.46) | 45.2 (4.58) | -2.98 (2.85) | -1.05 | 0.051 |
| SN | 39.4 (1.32) | 40.5 (1.41) | -1.13 (1.41) | -0.801 | 0.108 |

**Supplementary Table 5.** MAGIC LA-ICP-TOF-MS ^56^Fe concentration (ppm)

| Area | Control [<sup>56</sup>Fe] ($\sigma$) | MAGIC [<sup>56</sup>Fe] ($\sigma$) | $\Delta$[<sup>56</sup>Fe] ($\sigma$) | Z-score | p-value |
| --- | --- | --- | --- | --- | --- |
| CP | 87.6 (10.6) | 131 (24.5) | -43.9 (13.9) | -3.16 | 0.032 |
| GP | 72.6 (17.4) | 89.9 (14.9) | -17.3 (10.2) | -1.70 | 0.098 |
| SN | 81.3 (7.87) | 95.7 (13.0) | -14.4 (5.48) | -2.63 | 0.045 |

**Supplementary Table 6.**
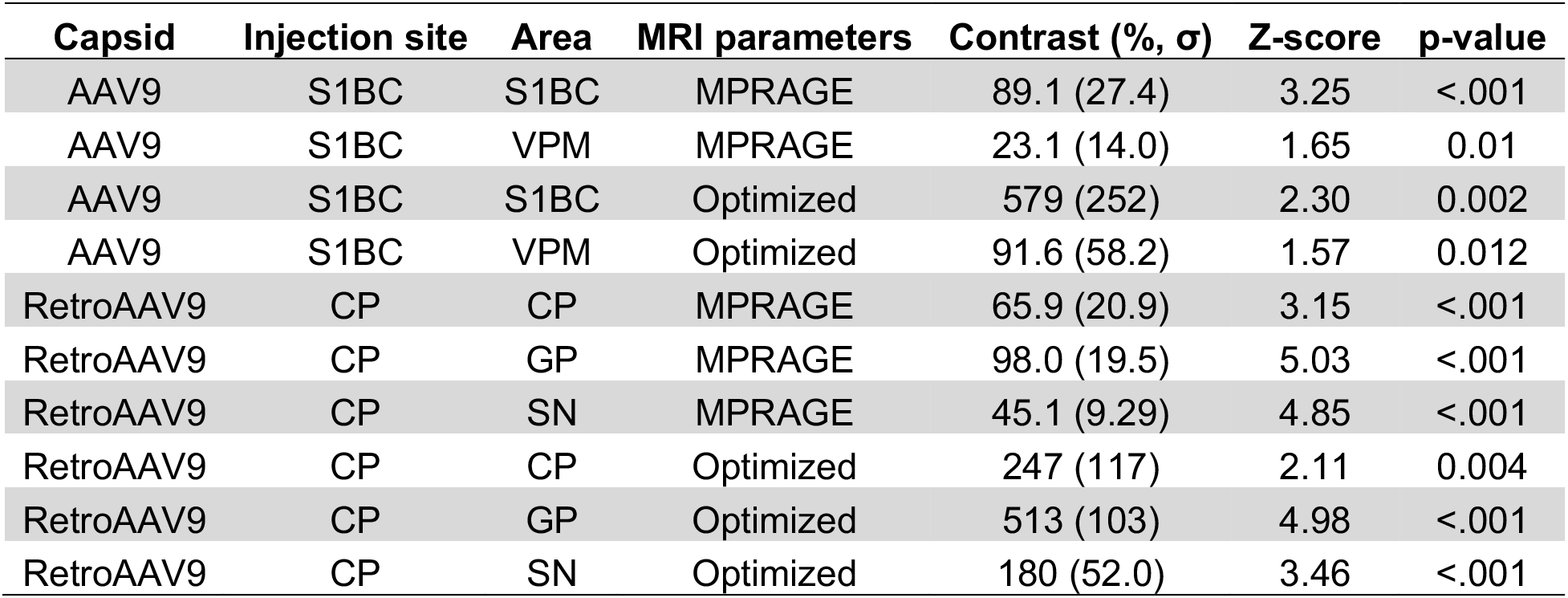
MAGIC+ effect size calculations.

| Capsid | Injection site | Area | MRI parameters | Contrast (%$\sigma$) | Z-score | p-value |
| --- | --- | --- | --- | --- | --- | --- |
| AAV9 | S1BC | S1BC | MPRAGE | 89.1 (27.4) | 3.25 | <.001 |
| AAV9 | S1BC | VPM | MPRAGE | 23.1 (14.0) | 1.65 | 0.01 |
| AAV9 | S1BC | S1BC | Optimized | 579 (252) | 2.30 | 0.002 |
| AAV9 | S1BC | VPM | Optimized | 91.6 (58.2) | 1.57 | 0.012 |
| RetroAAV9 | CP | CP | MPRAGE | 65.9 (20.9) | 3.15 | <.001 |
| RetroAAV9 | CP | GP | MPRAGE | 98.0 (19.5) | 5.03 | <.001 |
| RetroAAV9 | CP | SN | MPRAGE | 45.1 (9.29) | 4.85 | <.001 |
| RetroAAV9 | CP | CP | Optimized | 247 (117) | 2.11 | 0.004 |
| RetroAAV9 | CP | GP | Optimized | 513 (103) | 4.98 | <.001 |
| RetroAAV9 | CP | SN | Optimized | 180 (52.0) | 3.46 | <.001 |

## Supplementary Figures

**Supplementary Figure 1.**
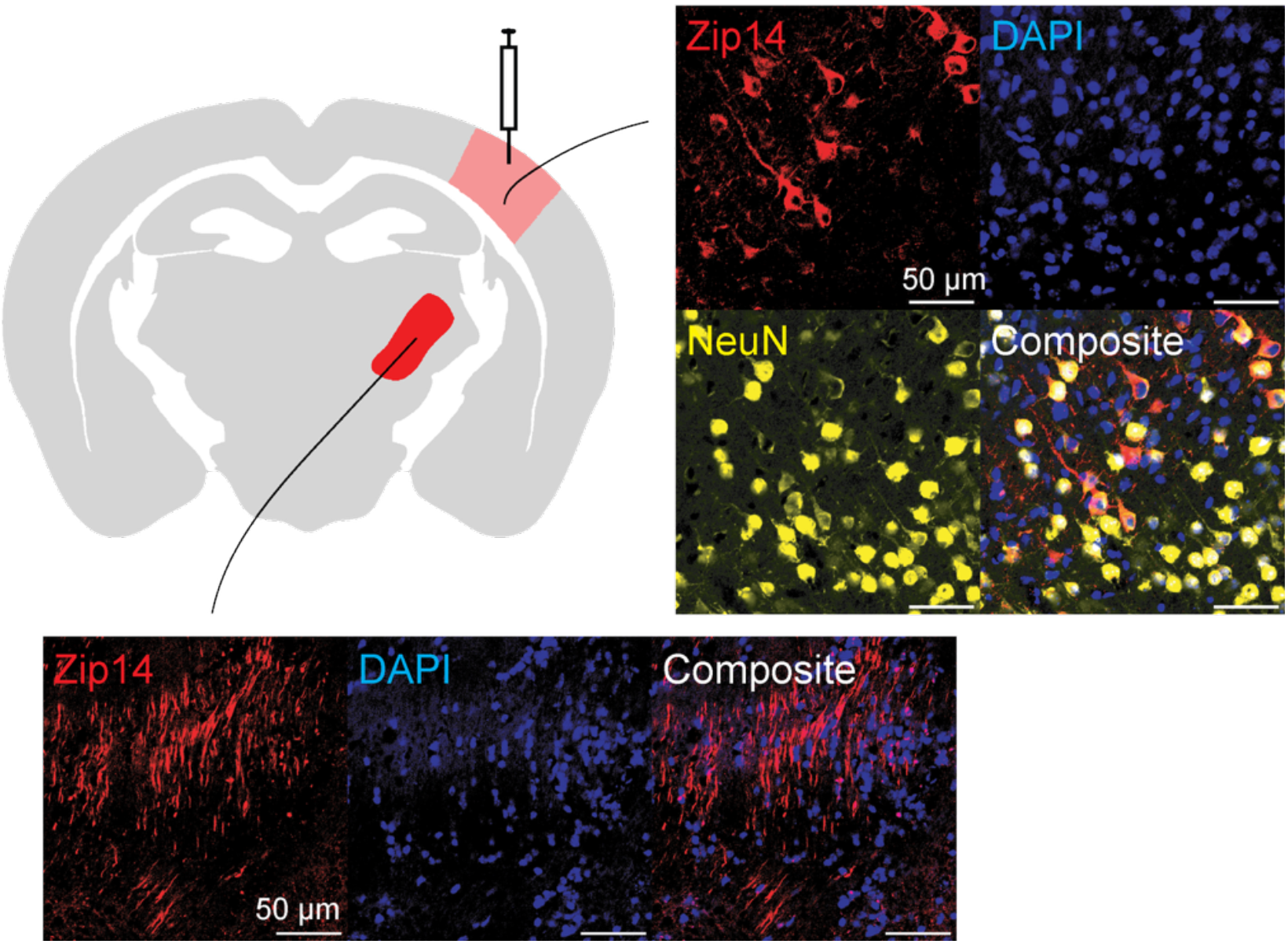
Immunofluorescence validation of Zip14 expression in S1BC neurons Matched immunofluorescence histology for Zip14 (red), DAPI (blue), and the neuronal marker NeuN (yellow). In the AAV9-Zip14 injected S1BC, Zip14 and NeuN double-positive cell bodies were detected. In the ipsilateral VPM, Zip14 positive axons and presynaptic terminals were detected.

**Supplementary Figure 2.**
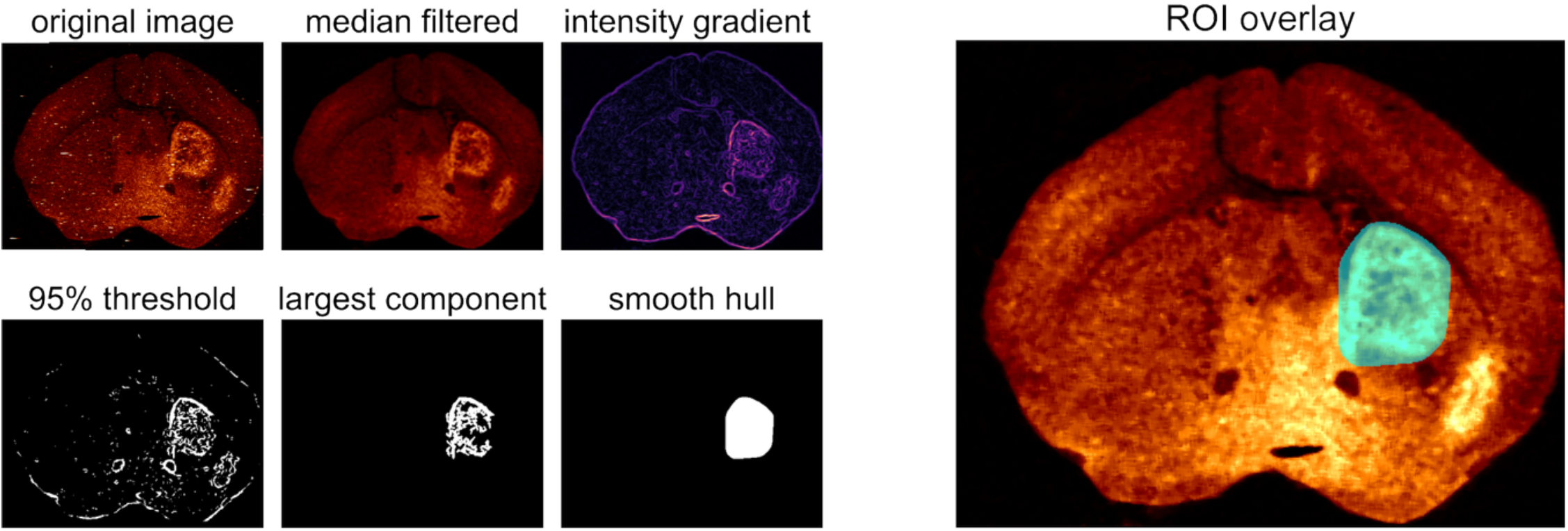
Automated detection of anomalous metal concentrations on LA-ICP-TOF-MS maps LA-ICP-TOF-MS derived ^55^Mn maps were median filtered to remove streaking artifact, and the magnitude intensity gradient was taken to detect edges. A 95% threshold was set, and the largest connected component was selected to exclude areas with smaller ^55^Mn increases. A continuous, closed ROI was defined by the smooth hull around the selected component.

**Supplementary Figure 3.**
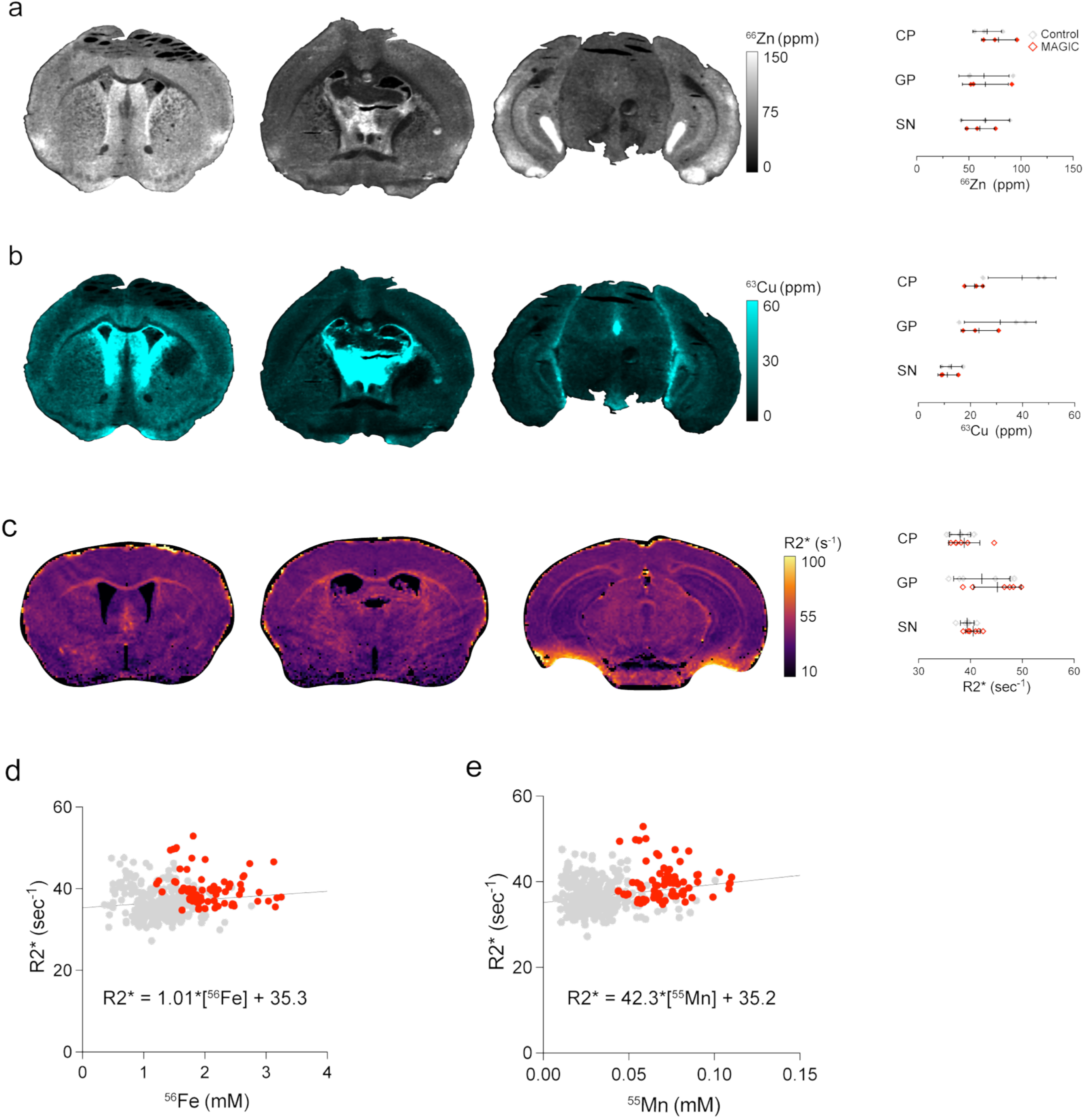
LA-ICP-TOF-MS measurements of Zn^2+^ and Cu^2+^, and R_2_^*^ correlation (a) LA-ICP-TOF-MS derived ^66^Zn maps and automated ROI-based quantification show no significant differences between Zip14 expressing and contralateral control areas. (b) Similar analysis of ^63^Cu maps showed a significant decrease in ^63^Cu in the injected CP compared to contralateral control, and no significant differences in the GP nor SN. (c) Representative R_2_^*^ mapping results reveal no observable R_2_* changes in any brain area. ROI-based quantification confirmed no significant differences in R_2_^*^ in any connected brain area. (d) Matched ROI measurements (n = 450) made on R_2_^*^ maps and ^56^Fe maps enabled measurement of in vivo transverse relaxivity for Fe^2+^. (e) Matched ROI measurements (n = 450) made on R_2_^*^ maps and Mn^55^ maps enabled measurement of in vivo transverse relaxivity for Mn^2+^. The measured r_2_^*^ for Mn^2+^ more closely matched literature reported values than that for Fe^2+^.

**Supplementary Figure 4.**
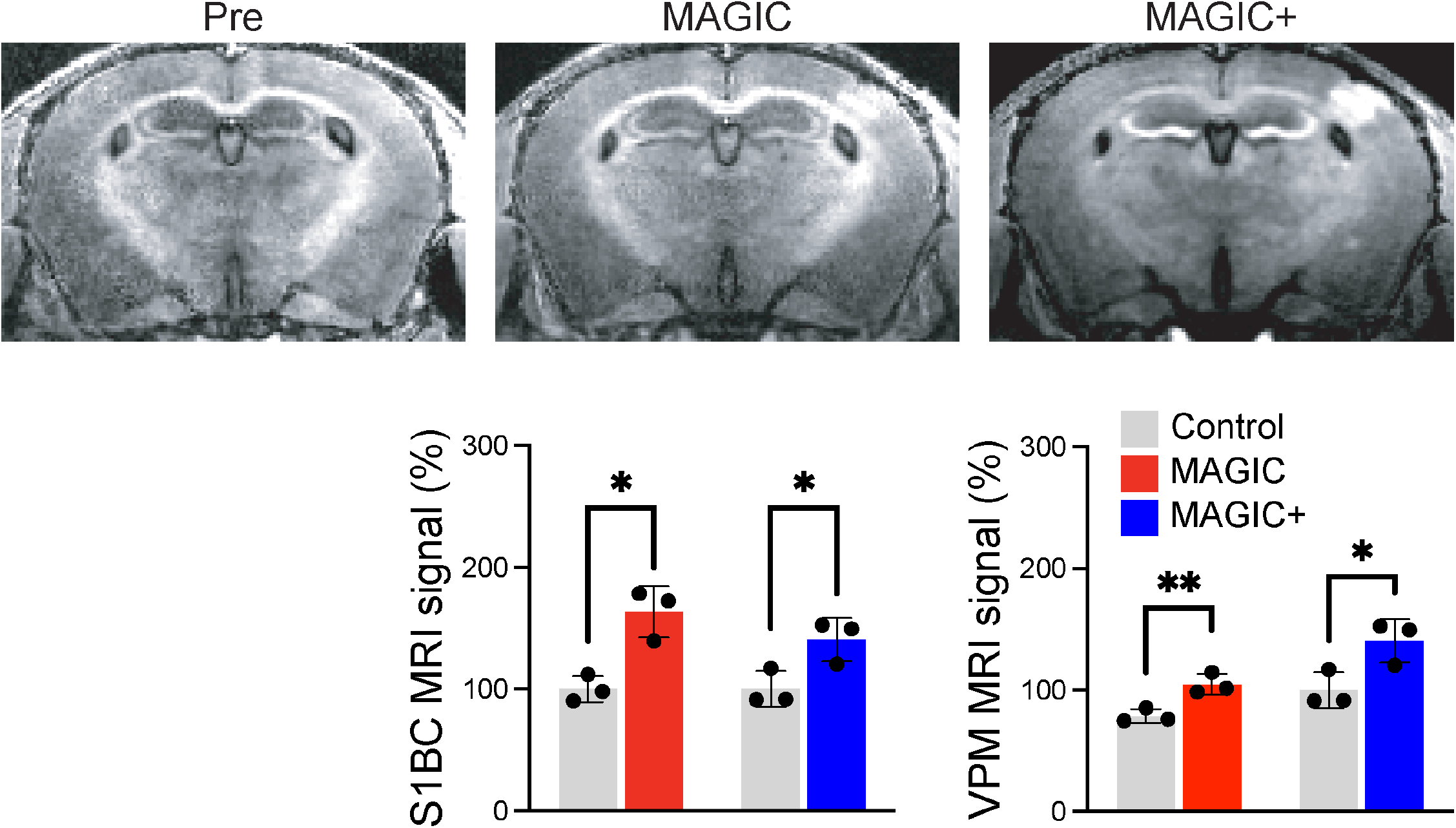
Half-dose MnCl_2_ supplementation (30 mg/kg) produces comparable MAGIC+ effects To test the effectiveness of lower supplemental MnCl_2_ doses to produce MAGIC+, 100 mM MnCl_2_ was delivered intraperitoneally to mice injected with AAV9-Zip14 into the S1BC at a dose of 30 mg/kg (n = 3). Representative Pre and MAGIC images matched previous results. Representative MAGIC+ images from the same animal after 30 mg/kg MnCl_2_ supplementation also closely matched previous results. Quantification of these effects revealed statistically significant increases in MAGIC and MAGIC+ in the injected S1BC and ipsilateral VPM. The magnitude of MAGIC+ was roughly in proportion to the supplemented MnCl_2_ dose. This result indicates that lower doses of supplemental MnCl_2_ have the potential to contribute to MAGIC+.

