## Supplementary material for "Manganese Accumulation for Genetically Induced Contrast (MAGIC) MRI in the brain across species": LA-ICP-TOF-MS supplementary methods

*Materials*

All standards, samples, and blanks were prepared gravimetrically using sub boiling point distilled (Savillex DST-1000, Eden Prairie, MN, USA) trace metal grade nitric acid (70%, Fisher chemical, Cat#: A509P212 for filter and gelatin digestions), ultrapure water (18.2 MΩ·cm @ 25 °C), and metal free polypropylene conical tubes (15 and 50 mL, Labcon, Petaluma, CA, USA). Gelatin standards were prepared using 300 bloom porcine gelatin (Sigma Aldrich, Cat# G2500, St. Louis, MO, USA) and a 1000 µg/mL IV-Stock-4 multi-element standard (Inorganic Ventures, Christiansburg, VA, USA) containing Ca, K, Ba, B, Cr, Cu, In, Pb, Mn, Ag, Sr, Zn, Mg, Al, Bi, Cd, Co, Ga, Fe, Li, Ni, Na, Tl and sodium phosphate monobasic (Sigma-Aldrich, Cat#: S0751, St Louis,MO, USA).

*Custom Gelatin Standard Preparation*

To prepare a homogeneous 10% (m/v) gelatin calibration standards with spiked multi-element standard, 1 g of gelatin was weighed using a plastic spatula onto weigh paper and slowly added to 8.95 mL of pre-warmed (55 °C) ultrapure water and mixed gently using a plastic spatula. The gelatin-water mixture is inverted gently to minimize air bubbles and degassed in an ultrasonic water bath set at 55 °C to remove any air bubbles during mixing. After the gelatin has mixed completely (no evidence of viscosity changes upon tube inversion), Either 50 µL of multi-element IV-Stock-4 standard is added and mixed gently resulting in a 50 µg/g high standard concentration or sodium phosphate monobasic is added to make a 4000 µg/g high NaP standard. High stock gelatin standards were then diluted into fresh 10% (w/v) gelatin to produce 25, and 12.5 µg/g of multi-element standard (50, 25, and 12.5 µg/g) and 2000 and 1000 µg/g of NaP standard (4000, 2000, 1000 µg/g) as well as blank gelatin for 4 points per calibration curve (50, 25, 12.5, 0 µg/g). 50 µL of the standard-gelatin mixtures were placed in individual pre-weighed 15 mL metal-free conical tubes, allowed to cool at room temperature, weighed and acid digested to determine concentrations via ICP-OES and ICP-QQQ-MS (see below for more details).

*Sample Preparation for Inductively Coupled Plasma (ICP) Analysis*

After preparing liquid gelatin standards and blanks, 50 µL of each standard was added to a pre-weighed 15 mL metal free conical tube and capped as to avoid evaporation of the heated liquid. After allowing the gelatin to cool (at least 4 hours at room temperature), 300 µL of distilled trace metal grade 70% nitric acid was added to each tube and digested at 70 °C for 4 hours. Following digestion, 9.7 mL of ultrapure deionized H_2_O was added, and the sample was weighed.

*ICP-Optical Emission Spectroscopy (OES) analysis of Gelatin Standards*

NaP gelatin standards were analyzed using an Agilent 5800 ICP-OES (Agilent, Santa Clara, CA, USA) equipped with the Agilent SPS 4 Autosampler, quartz cyclonic spray chamber, micromist nebulizer, and AVS 6/7 valve introduction system. Daily instrument performance is validated via a detector and multiwavelength calibration in axial and radial mode using the manufacturer’s multi-element calibration standard containing 50 mg/L Al, As, Ba, Cd, Co, Cr, Cu, Mn, Mo, Ni, Pb, Se, Sr, Zn and 500 mg/L K in 5% nitric acid. ICP-OES standards were prepared from individual stock solutions of Na and P standards (Inorganic Ventures, Christiansburg, VA, USA) that were diluted with 3% (v/v) trace nitric acid in ultrapure deionized water to a final element concentration of 100, 50, 25, 12.5, 6.25, 3.125, and 0 (blank) µg/g standard. Internal standardization was accomplished inline using the AVS 6/7 valve and a 1 µg/g internal standard solution in 3% (v/v) trace nitric acid in ultrapure water consisting of Y (Inorganic Ventures, Christiansburg, VA, USA). The emission lines selected (in both axial and radial mode) for analysis were Na (589.592 nm), P (213.618 nm), and Y (371.029 nm) used for internal standardization.

*ICP Triple Quadrupole Mass Spectrometry (ICP-QQQ-MS) Analysis of Gelatin Standards*

Multi-element gelatin standards were analyzed using an Agilent 8900 Triple Quadrupole ICP-MS (Agilent, Santa Clara, CA, USA) equipped with the Agilent SPS 4 Autosampler, integrate sample introduction system (ISiS), x-lens, and micromist nebulizer. Daily tuning of the instrument was accomplished using the manufacturer supplied tuning solution containing Li, Co, Y, Ce, and Tl. Global tune optimization was based on optimizing intensities for ^7^Li, ^89^Y, and ^205^Tl while minimizing oxides (^140^Ce^16^O/^140^Ce < 1.5%) and doubly charged species (^140^Ce++/^140^Ce+ < 2%). Following global instrument tuning, gas mode tuning in He KED and O_2_ mode was accomplished using the same manufacturer supplied tuning solution. In KED mode (using 100% UHP He, Airgas), intensities for ^59^Co, ^89^Y, and ^205^Tl were maximized while minimizing oxides (^140^Ce^16^O/^140^Ce < 0.5%) and doubly charged species (^140^Ce++/^140^Ce+ < 1.5%) with short term RSDs < 3.5%. In O_2_ mode (using 100% UHP O_2_, Airgas) intensities for ^59^Co, ^89^Y, and ^205^Tl with short term RSDs < 3.5%. ICP-MS standards were prepared from a stock solution of IV-Stock-4 multi-element standard (As, B, Ca, Cd, Co, Cr, Cu, Fe, K, Mg, Mn, Mo, Na, Ni, Pb, S, Se, V, and Zn, Inorganic Ventures, Christiansburg, VA, USA) that were diluted with 3% (v/v) trace nitric acid in ultrapure deionized water to a final element concentration of 100, 50, 25, 12.5, 6.25, 3.125, 1.5625, and 0 (blank) ng/g standard. Internal standardization was accomplished inline using the ISIS valve and a 200 ng/g internal standard solution in 3% (v/v) trace nitric acid in ultrapure water consisting of Bi, In, ^6^Li, Sc, Tb, and Y (IV-ICPMS-71D, Inorganic Ventures, Christiansburg, VA, USA). The isotopes selected for analysis were ^39^K, ^55^Mn, ^56^Fe, ^59^Co, ^60^Ni, ^65^Cu, and ^66^Zn, with ^6^Li, ^45^Sc, and ^89^Y used for internal standardization.

*Gelatin Standard Sectioning Protocol*

Gelatin was prepared for cryosectioning using a Leica CM 3050 S (Leica Biosystems, Deer Park, IL, USA) cryostat. After the gelatin is dissolved and the solution is still at 55 °C, approximately 300 µL of warmed gelatin standard is pipetted onto a precooled chuck inside the cryostat (-20 °C). Gelatin is allowed to freeze for no less than 4 minutes at -20 °C. Once the gelatin standards were completely frozen, samples were sectioned using a -21 °C chamber temperature and -20 °C objective temperature (temperatures were adjusted slightly depending on the quality of the sections and whether the section stuck to the blade or anti-roll plate) and a section thickness of 20 µm to match the section thickness of the tissue. Gelatin sections were then transferred to pre-cleaned charged microscope slides (Superfrost plus, thermos fisher scientific) and kept at

-20 °C until laser ablation.

*LA-ICP-TOF-MS instrument setup and sample parameters*

Glass slides were loaded into a Bioimage 266 nm laser ablation system (Elemental Scientific Lasers, Bozeman, MT, USA) which is equipped with an ultra-fast low dispersion TwoVol3 ablation chamber and a dual concentric injector (DCI3) and is coupled to an icpTOF S2 (TOFWERK AG, Thun, Switzerland) ICP-TOF-MS. Daily tuning of the LA-ICP-TOF-MS settings was performed using NIST SRM612 glass certified reference material (National Institute for Standards and Technology, Gaithersburg, MD, USA). Optimization for torch alignment, lens voltages, and nebulizer gas flow was based on high intensities for ^140^Ce and ^55^Mn while maintaining low oxide formation based on the ^232^Th^16^O+/^232^Th+ ratio (< 0.5). A list of instrument parameters for LA-ICP-TOF-MS is summarized in Table S1.

In short, 20 µm thick gelatin standards (3 multi-element standards, 3 NaP standards, and a blank) were placed directly on charged slides or Kapton tape which was then placed sticky side up on charged slides. These 2 slides were loaded with a sample slide on the LA sample holder and loaded into the LA system. The system is purged for 5 minutes and patterns are selected on the standards and samples of interest. For gelatin standards, 10 lines going across each standard and the blank were drawn using the same laser parameters as the samples (see table). Specifically, 20 µm circular laser spot sizes at 70% laser power and 100 Hz repetition rate were used with an interline distance 3 times greater than the spot size. This allowed for clean ablation of individual lines without overlap. For the sample, reference points were made around the sample and the aforementioned laser parameters were used with an interline distance equal to the laser spot (20 µm) with no overlap to sample the entire tissue.

*Data Acquisition and Analysis of LA-ICP-TOF-MS Data*

Data was recorded using TofPilot 1.3.4.0 (TOFWERK AG, Thun, Switzerland). The LA-ICP-TOFMS data were saved in the open-source hierarchical data format (HDF5). Post-acquisition data processing was performed using our in house developed AutoSpect software pipeline (Crawford, A. M., Zee, D. Z., Jin, Q., Sue, A., Sinha, N., Ahn, S. H., O’Halloran, T. V. & MacRenaris, K. W. AutoSpect: an all-in-one software solution for automated processing of LA-ICP-TOF-MS datasets. Journal of Analytical Atomic Spectrometry 40, 2162–2178 (2025).). The data processing comprised the following steps: (1) drift correction of the mass peak position in the spectra over time via time-dependent mass calibration, (2) determining the peak shape, and (3) fitting and subtracting the mass spectral baseline. The data was further processed with Iolite version 4.8.6 (Elemental Scientific Lasers, Bozeman, MT, USA). For calibration, signal responses for each ablation line per gelatin standard were fit to a linear regression and spline auto smoothed. Calibration curves are then generated and using 3D trace elements inside of DRS in Iolite, we can convert integrated counts per second to µg/g (using the previous calculations for the gelatin standards via ICP). Representative calibration curves are shown below.

Supporting Information - Keith

**Table S1.** ICP-TOF-MS and laser ablation parameters for LA-ICP-TOF-MS analysis of tissues

| *ICP-MS Parameters (Tofwerk S2)* | | |
| --- | --- | --- |
| Parameter | **Unit** | **Value** |
| RF Power | W | 1550 |
| Sampling Depth | mm | 5 |
| Cone Material | Nickel | - |
| Cone Insert (STD) | mm | 3.5 |
| Plasma Gas Flow | L/min | 14.0 |
| Auxillary Gas Flow | L/min | 8.0 |
| Nebulizer Gas Flow | L/min | 0.95-1.02 |
| Measurement Mode | CCT Mode | - |
| CCT Gas Flow (100% He) | mL/min | 5 |
| CCT Focus lens | V | -10.09 |
| CCT Entry Lens | V | -191.31 |
| CCT Mass | V | 246 |
| CCT Bias | V | -4 |
| CCT Exit Lens | V | -173 |
| *Time-of-Flight Parameters (Tofwerk S2)* | | |
| Parameter | **Unit** | **Value** |
| m/z range | amu | 14 - 256 |
| Resolution | m/Δm | 1000 |
| ODG Settings | ms | 10 |
| Notch Calibration |  | .878 |
| Notch (40 amu) | V | 1.5 |
| Notch (28 amu) | V | 1.2 |
| Notch (35.5 amu) | V | 1.6 |
| Notch (15.9 amu) | V | 2.6 |
| *Laser Ablation Parameters (ESL Bioimage 266 nm)* | | |
| Parameter | **Unit** | **Value** |
| Spot Size | µm | 20 |
| Interline Distance (y) | µm | 20 |
| Overlap (x) | µm | 0 |
| Repetition Rate | Hz | 100 |
| Laser Power | % | 70 |
| Laser Fluence | J/cm^2^ | 1.7-2.1 |
| Sample Energy | mJ |  |
| Imaging Cup Flow Rate (He) | mL/min | 350 |
| Imaging Chamber Flow Rate (He) | mL/min | 300 |
| PEEK Tubing I.D. | mm | 1 |

**Supporting Figure 1.** ^56^Fe 3D surface, Block linear curve fit, and multiple fits (10 standard lines)

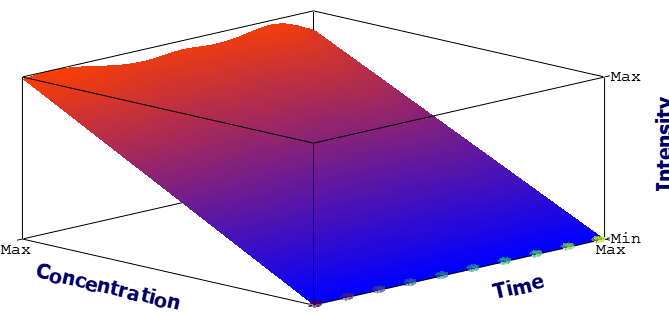

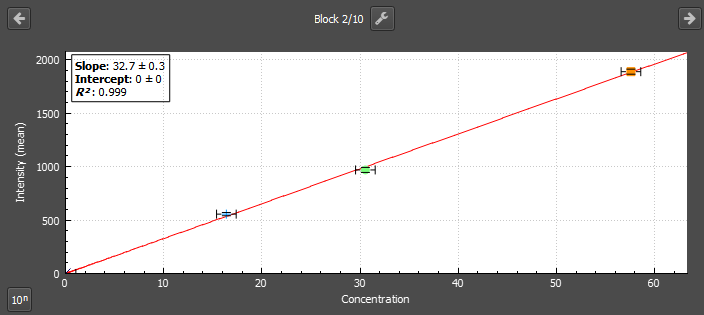

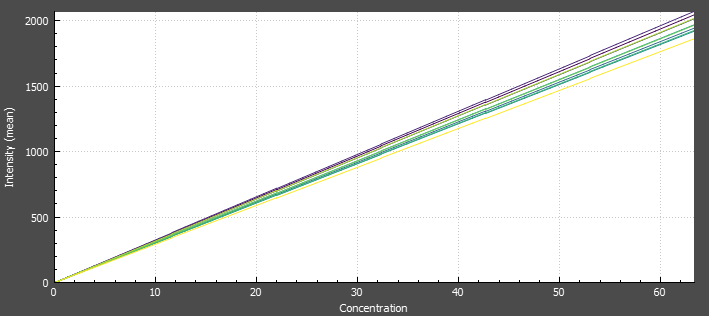

**Supporting Figure 1.** ^55^Mn 3D surface, Block linear curve fit, and multiple fits (10 standard lines)

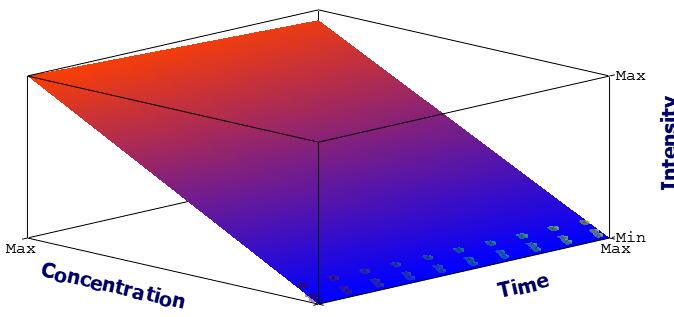

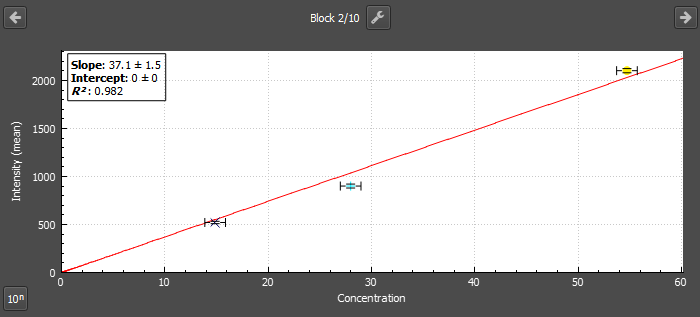

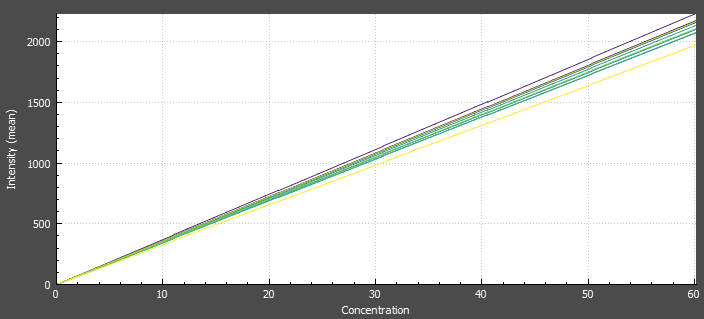
